# DET1 dynamics underlie co-operative ubiquitination by CRL4^DET1^-COP1 complexes

**DOI:** 10.1101/2024.06.08.598067

**Authors:** Abigail E Burgess, Tarren A Loughran, Liam S Turk, Jess L Dunlop, Sam A Jamieson, Jack R Curry, Pavel Filipcik, Simon HJ Brown, Peter D Mace

## Abstract

Ubiquitin ligases regulate core cellular processes through diverse mechanisms. The ubiquitin ligase COP1 is conserved from plants to humans and is particularly important for targeting developmental transcription factors for ubiquitination. COP1 can function independently, but can also be recruited to Cullin-4 ubiquitin ligase complexes via the DET1 adaptor protein. However, the mechanism of action of complexes containing COP1 and DET1 is not well understood. Here we report the cryo-electron microscopy structure of human DET1, bound to proteins that enable Cullin-4 recruitment (DDB1-DDA1) and an additional ubiquitin ligase enzyme (Ube2e2). We observe that DDA1 stabilises a closed conformation of DET1, binding adjacent to a unique Ube2e2 binding-insert in DET1. Moreover, we demonstrate that closure of DET1 underlies COP1 recruitment, which binds in an antiparallel dimeric state. Disrupting either the Ube2e2-binding insertion of DET1, or distinct recruitment sites on COP1, abolish DET1–COP1 binding and DET1-mediated modulation of COP1 levels. The multifaceted architecture provides an efficient platform for ubiquitination of substrates, or COP1 itself, by Cullin-4^DET1^ and offers multiple opportunities for physiological regulation.

## INTRODUCTION

Ubiquitin-mediated post-translational modification is a pervasive biological signaling mechanism. It encompasses modifications ranging from a single ubiquitin attachment to chains linked at lysine branch points, governing diverse outcomes for modified proteins. Chains attached via lysine 48 or lysine 11 facilitate proteasomal degradation of modified proteins^1,2^. One important role of degradative ubiquitination is regulating the levels of transcription factor; attaching degradative ubiquitin chains targets specific transcription factors for proteasomal degradation, thereby facilitating the remodeling of gene expression patterns. Transcriptional regulation via ubiquitin involves a variety of ligases, including standalone ubiquitin ligases and multimeric complexes with interchangeable modules, enabling the regulation of different substrate sets.

Constitutive Photomorphogenic 1 (COP1) is a ubiquitin ligase that is essentially conserved from plants to humans. The main targets of COP1 are transcriptional regulators, and COP1-mediated ubiquitination acts a crucial step in defining differentiation and developmental patterning. COP1 was first identified in plants as a preeminent regulator of light-mediated development^3^, where COP1-based ubiquitination is controlled by photoreceptors and environmental stimuli^4,5^. Human COP1 is not light-regulated but shares the same core function, it is involved in the ubiquitination of a range of key developmental transcription factors—including p53, c-Jun, and ETS and C/EBP families^6–10^. Through its ubiquitin ligase activity, COP1 has been shown to regulate immune, glial, hepatic, and pancreatic cell function, and be relevant to traditional cancer treatments based on kinase inhibition and immune checkpoint blockade therapy^11–16^.

Both plant and human COP1 are known to be able to function with another protein, De-etiolated-1 (DET1), to degrade some substrates^3,9,17^. DET1 has a well-established role as a recruitment factor for the Cullin-4 (CUL4) ubiquitin ligase (CRL4), a major nuclear platform for building ubiquitin chains in partnership with the E3 ligase Rbx1^18–21^. For CRL4, substrate adapters like DET1 are recruited through the DNA Damage binding Protein 1 (DDB1).

Exchanging the adaptor switches substrate specificity and other related DCAF (DDB1 and CUL4-associated factor) proteins also bind to CUL4^22^. Regulation of CRL4 activity comes at several levels, including modification by the ubiquitin-like molecule NEDD8 and exchange of DCAF proteins^23–25^. Accordingly, many DCAF proteins have received significant attention as recruitment modules in the field of targeted protein degradation^26^.

Like many DDB1 partners DET1-DDB1 also associates with an accessory protein called DDA1^27^, and the resulting complex can exist in isolation and has been denoted as the ‘DDD’ (DDB1-DDA1-DET1) complex^3,28^.

DET1 is unique amongst DCAF proteins in that it can directly bind ubiquitin conjugating (E2) enzymes. When part of the DDD complex, DET1 can specifically the human Ube2e family, through an ill-defined mechanism^28^. DET1 in plants binds to a catalytically inactive ubiquitin conjugating enzyme called COP10, suggesting an evolutionarily conserved function that may not depend on its catalytic activity^29,30^. The exact role played by the E2s recruited by DET1 is currently unclear. There have been few direct substrates reported to bind human DET1, and so it appears the clearest role of human DET1 is in binding to COP1. Overall, DET1 appears to occupy a unique juncture amongst the DCAFs—by associating with both COP1 and CRL4, it potentially acts as a link between two major ubiquitin ligase complexes.

The domain structure of COP1 gives it capacity to both recruit substrates and promote ubiquitin transfer—containing an N-terminal RING domain, a central coiled-coil region, and a C-terminal WD40 domain primarily responsible for substrate recruitment^31^. However, COP1 appears to have multiple modes of activity: ubiquitinating some substrates bound to its WD40 domain with its own RING domain; and modifying other substrates independently of its RING domain when COP1 associates with CRL4^DET1^ complexes. Binding between COP1 and DET1 is known to be regulated by ERK-family MAP kinase phosphorylation^10^, and mutations in either DET1 or COP1 in cancer can drive resistance to ERK-pathway targeted therapies^16^. Complexes formed by DET1 and COP1 clearly play a key role in the developmental programming within normal cells, disease progression, and therapeutic resistance. However, the architecture of complexes that contain both CRL4 and DET1, with or without COP1, and contributions of accessory proteins such as DDA1 and additional ubiquitin-conjugating enzymes to larger complexes is currently unclear.

Here we use a combination of structural and biochemical methods to delineate the architecture and mechanism of DET1-COP1 complex assembly. We use cryo-electron microscopy to determine the high-resolution structure of the DDB1-DDA1-DET1-Ube2e2 complex, which clearly defines a novel binding site for Ube2e2 and explains subclass specificity of DET1-E2 binding. Structural variability analyses show that DET1 exists in a range of dynamic states—from relatively open, to a closed state stabilised by DDA1 when Ube2e2 is bound. AlphaFold2 modelling suggests that the closed state of DET1 is essential for recruitment of an anti-parallel COP1 homodimer. Cellular degradation experiments show that E2 recruitment and COP1 binding interfaces—both of which require DET1 closure— each contributing to assembly of the functional CRL4^DET1^-COP1 ligase.

## RESULTS

### Architecture of the human DDD–Ube2e2 complex

To gain insight into the interactions and function of the DDD complex we established a system to co-express and purify the human DDD complex with full-length Ube2e2 from insect cells. Following affinity purification, subsequent purification by size-exclusion chromatography showed a significant proportion of Ube2e2 was retained in a high molecular weight complex with core DDD components (Figure 1b). Full-length DDB1, DET1, DDA1 and Ube2e2 were used for initial structure determination by cryo-EM, which resolved all four components but showed severe particle orientation bias and map anisotropy (Supplementary Figure 1). We subsequently purified the DDD complex with DDB1 lacking its central WD40 (BPB^32^) domain, which can exhibit significant flexibility^33^. The BPB domain would normally bind to CUL4 but has been deleted to facilitate high-resolution structures of other DCAF proteins^34,35^. Eventual maps of DDD(ΔBPB)-Ube2e2 markedly improved map quality and allowed modelling of continuous density for most of the complex to an FSC-determined resolution of 2.84 Å (Figure 1c; Table 1; Supplementary Figure 2).

**Figure 1:**
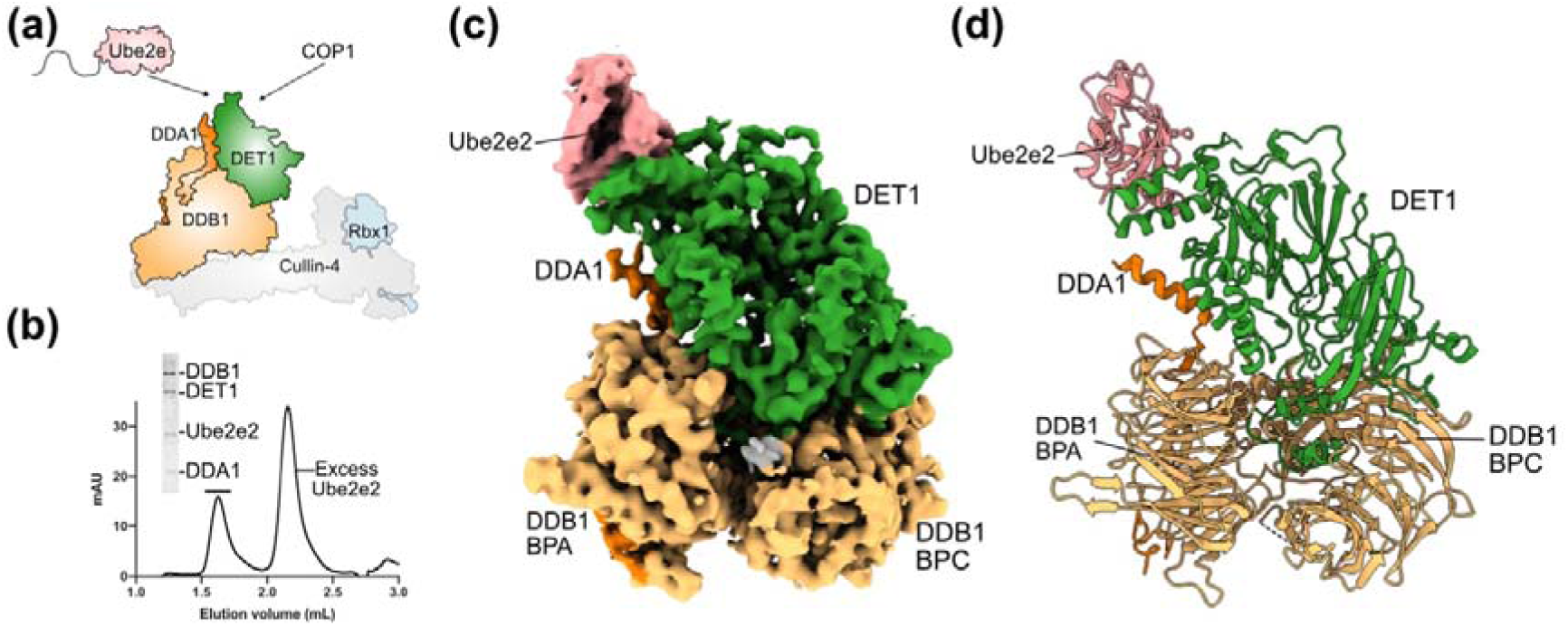
Structure of the DDD:Ube2e2 complex. **(a)** Schematic representation of the DDD complex. **(b)** Purification of Ube2e2 with the DDD complex. DDD complex was purified using size exclusion with an excess of Ube2e2. **(c)** Cryo-EM density of the DDB1(ΔBPB)-DET1-DDA1-Ube2e2 complex. **(d)** Molecular model of DDB1(ΔBPB)-DET1-DDA1-Ube2e2 complex. DET1 (green), DDB1(ΔBPB) (yellow), DDA1 (orange), Ube2e2 (pink).

**Table 1:** Cryo-EM data collection, refinement, and validation statistics.

| **Data Collection and Processing** | **DDD(ΔBPB)**<br>**Complex** | **Cluster 1** |
| --- | --- | --- |
| Magnification | 130000x | 130000x |
| Voltage (keV) | 300 | 300 |
| Electron exposure ($e^-/\text{\AA}^2$) | 80 | 80 |
| Defocus ($\mu\text{m}$) | -1.0 | -1.0 |
| Pixel size ($\text{\AA}$) | 0.65 | 0.65 |
| Symmetry imposed | C1 | C1 |
| Final particle images (no.) | 61234 | 70496 |
| Map resolution fourier shell correlation (FSC) threshold | 0.143 | 0.143 |
| Map resolution ($\text{\AA}$) | 2.83 | 3.02 |
| **Refinement** |  |  |
| Initial model used (PDB) | AlphaFold |  |
| Model resolution FSC threshold ($\text{\AA}$) | 0.5 | |
| Model resolution ($\text{\AA}$) | 3.4 | |
| Model composition |  |  |
| Non-hydrogen Atoms | 11999 |  |
| Protein residues | 1527 |  |
| B-factor ($\text{\AA}^2$) | | |
| Minimum | 70.82 |  |
| Maximum | 281.24 |  |
| Mean | 154.54 |  |
| RMS deviations |  |  |
| Bond lengths ($\text{\AA}$) | 0.002 | |
| Bond angles ($^\circ$) | 0.531 | |
| Validation |  |  |
| MolProbity score | 1.63 |  |
| Clashscore | 7.89 |  |
| Rotamer outliers (%) | 0.15 |  |
| Ramachandran plot |  |  |
| Favoured (%) | 96.75 |  |
| Allowed (%) | 3.25 |  |
| Outliers (%) | 0 |  |

The overall model of the DDD(ΔBPB)-Ube2e2 complex shows the N-terminal H-box motif of DET1 interacting with the BPC of DDB1, as expected based on comparable DCAF-DDB1 structures^22^. While DET1 binds in the same pocket of DDB1, there are some subtle differences (Supplementary Figure 3). The arrangement of DET1 in the BPC pocket is most similar to DCAF1 and DCAF15, binding via a helix-loop-helix motif. Comparing more widely shows that this mode of interaction is relatively minimal compared to interactions of Cereblon, DDB2, and DCAF16, which each contain additional elements mediating interactions with DDB1 (Supplementary Figure 3).

Beyond the N-terminal region, DET1 is comprised of a predominantly beta sheet structure, which overlays relatively well with the AlphaFold2 predicted structure of DET1 (RMSD of pruned atom pairs - 0.931 Å, across all 508 atom pairs – 1.288 Å, Supplementary Figure 4). The major departure of DET1 from its beta sheet core is an insertion of residues 262–329, which have relatively higher LDDT scores in the AlphaFold2 model of DET1. These residues form a series of alpha-helices propagating from the core at approximately right angles, having the effect of forming a ‘cup’, within which the core ubiquitin-conjugating domain of Ube2e2 is well-defined by cryoEM density. The extended structure of DDA1 wraps around DDB1— with residues 3-17 binding to the bottom of the BPA WD40 domain, and its C-terminal helix (residues 54–68) interacting below the E2 binding site of DET1 (Figure 1c and Supplementary Figure 4b). The segment connecting residues 17-42 of DDA1 is largely undefined by cryo-EM density.

### A DET1 ‘cup’ binds the E2-backside

There are two major components of the DET1 cup that interact with the C-terminal helix and core beta sheet of Ube2e2 respectively. A segment composed of residues 257–291 mediates interactions towards the C-terminus of the E2, primarily through a helix formed by residues 264–271 and Phe291 (Figure 2a). Residues 292–329 of DET1 form a helix-loop-helix arrangement that makes extensive contacts with the beta-sheet backside of Ube2e2 (Figure 2a). Comparing the sequence and AlphaFold predictions of the equivalent regions of human and Arabidopsis DET1 shows that the region equivalent to human 257–291 contains an extended helix and large insertion with low LDDT scores. In contrast, the helix-loop-helix is highly conserved—residues 314–358 of Arabidopsis DET1 are predicted to form a helix-loop-helix that would be well placed to bind to the COP10 ubiquitin conjugating enzyme that associate with Arabidopsis DET1 (Figure 2a)^29^. This suggests that the helix-loop-helix of DET1 is the core conserved element of E2 binding and specifically evolved to bind to the core of ubiquitin conjugating domain.

**Figure 2:**
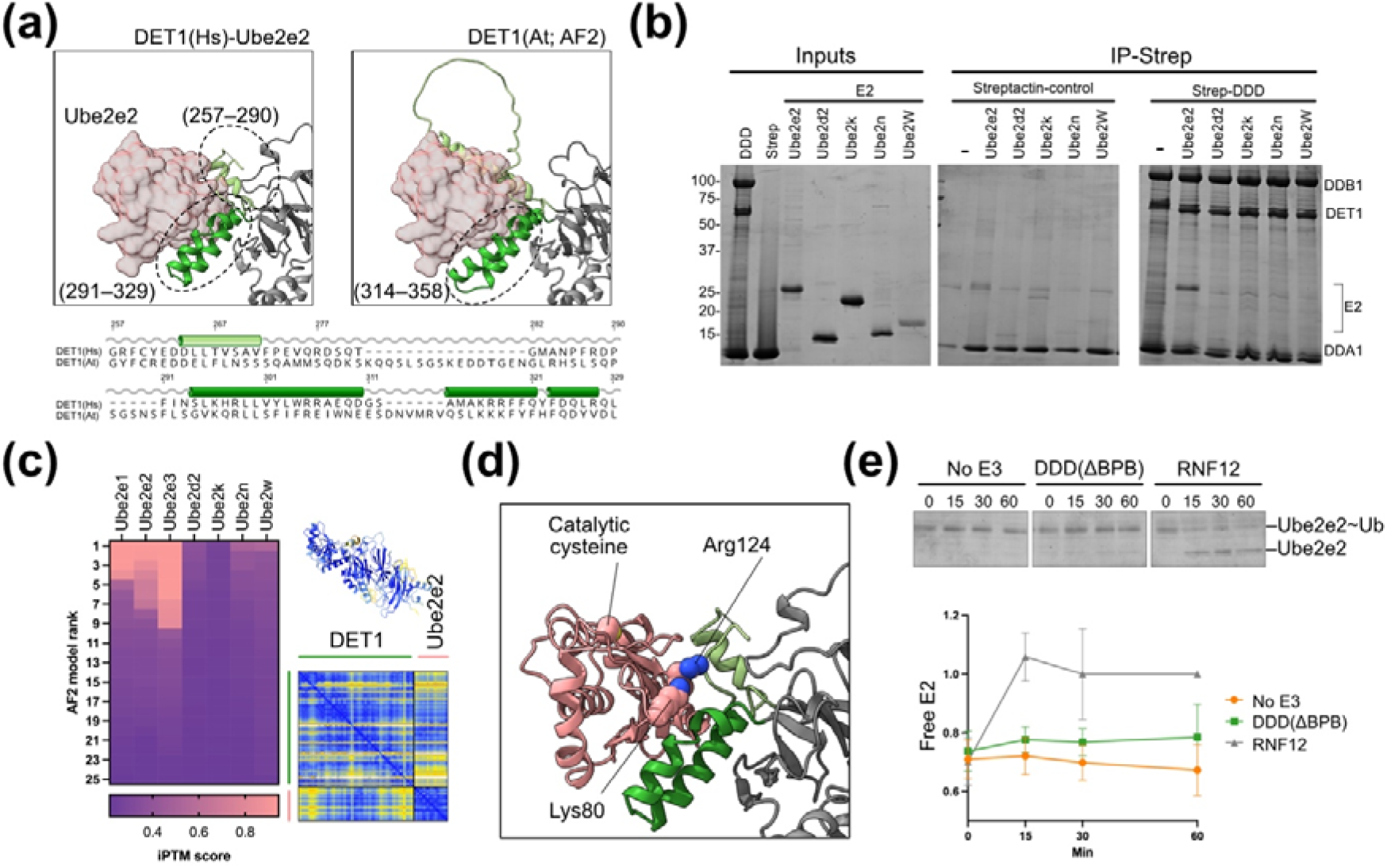
Interactions and specificity of E2 recruitment by DET1. **(a)** AlphaFold modelling and sequence alignment of the human and Arabidopsis E2-binding loop of DET1 shows conservation of a helix-loop-helix which mediates the interaction between DET1 and Ube2e2. (**b)** In vitro pull down of E2 binding to the DDD complex. Coomassie-stained gels showing pulldown of E2 proteins by the full-length DDD complex, immobilized on Strep-TactinXT resin, show that Ube2e2 alone binds the full-length DDD complex **(c)** AlphaFold modelling was used to screen DET1-E2 interactions. Predicated confidence of the interface is shown as a summary score-with only Ube2e family and COP10 having an IPTM score >0.8 across multiple models. PAE plot shows confidence in the DET1-Ube2e2 model. **(d)** Conserved residues in Ube2e family and COP10 drive specificity. Lys80 and Arg124 are conserved within the Ube2e family and COP10 but are not seen in any of the other E2s sampled. **(e)** DDD (ΔBPB) complex does not cause ubiquitin discharge from Ube2e2. Ubiquitin discharge assays were used to determine the ability of DDD to cause ubiquitin discharge from Ube2e2. Samples were taken at 0, 15, 30 and 60 mins and ubiquitin discharge was determined by staining gels with Coomassie Blue (left). Ube2e2 discharge was not enhanced by the presence of the DDD (ΔBPB) complex when compared to the no E3 control, indicating that the DDD (ΔBPB) complex does not cause ubiquitin discharge from Ube2e2.

Previous analysis of DET1 interactions with ubiquitin conjugating enzymes had suggested that the N-terminal extension of the of Ube2e family is required for DET1 binding, with some requirement for the core catalytic domain^28^. Our structure clearly places the ubiquitin conjugating domain but does not identify clear density for the N-terminus even though it was present in the samples used for cryo-EM sample preparation. While the previous work showed DET1 co-purifying with Ube2e1, Ube2e2, and Ube2e3^28^, clear interactions with the structurally conserved core catalytic domain raised the question of whether DET1 may also bind other ubiquitin conjugating enzymes beyond the Ube2e family.

To interrogate the specificity of the interaction between the Ube2e family and DET1, we compared the DET1 binding ability to a set of E2s. The E2s were chosen based on high sequence conservation to Ube2e2; including Ube2d2, Ube2n, Ube2k and Ube2w (Supplementary Figure 5). In vitro pulldown experiments with this set of E2s showed clear specificity—only Ube2e2 was pulled down by the DDD complex (Figure 2b). In parallel, we employed AlphaFold2 multimer modelling, where structures of full-length DET1 were predicted with the core-ubiquitin conjugating domains of the same set of E2 enzymes. In addition to those tested experimentally, our AlphaFold2 screen included human Ube2e1 and Ube2e3 with human DET1, along with the Arabidopsis COP10 protein, which is catalytically dead but associates with Arabidopsis DET1. AlphaFold2 screening showed remarkable DET1 specificity, with only human Ube2e1, Ube2e2, Ube2e3 and Arabidopsis COP10 showing iPTM scores >0.8 (Figure 2c), and high confidence predicted aligned error (PAE) plots.

Manual inspection of high-scoring structures (iPTM >0.80) revealed complexes that superimpose well into cryo-EM density, whereas structures with iPTM <0.80 showed E2 in the correct general area, but in a variety of different arrangements (Supplementary Figure 5a). This aligns well with initial observations that only the Ube2e family enzymes co-purify with the DDD complex^28^.

In combination, the experimental data and computational structures show that the interaction specificity is clearly driven by the ubiquitin conjugating domain, rather than by the N-terminal extension of E2 enzymes. To gain insight into residues that drive specificity, we analysed conservation of E2 residues that interact with the DET1 cup from the cryo-EM structure (Supplementary Figure 5b). This alignment showed that most interface residues are relatively conserved, apart from Lys80 and Arg124 (Ube2e2 numbering). Lys80 and Arg124 sit directly adjacent to the DET1 binding site (Figure 2d). This pair of basic residues are conserved in Ube2e1–3 and Arabidopsis COP10 but not in other closely related E2s. While residing at the DET1 interface, they do not directly overlap with the canonical backside ubiquitin binding site of E2 enzymes^36^. Thus, these residues appear to be well placed to differentiate DET1 binding but have an unclear impact on wider allosteric regulation by ubiquitin.

Binding to the beta-sheet core of Ube2e proteins is notable because of its overlap with the well-studied ‘backside’ ubiquitin-binding site^36,37^. Backside ubiquitin binding allosterically primes E2-Ub thioester cleavage from the active site cysteine catalysed by RING domain proteins^36,38,39^. While DET1 binding would clearly prevent backside ubiquitin binding, we hypothesised that DET1 cup residues may themselves promote E2-Ub conjugate discharge. This would explain previous observations that the majority of Ube2e family molecules associating with the DDD are in the uncharged, rather than ubiquitin-conjugated state^28^. To test this, we measured ubiquitin discharge from Ube2e2 in the presence of the complete DDD complex. Discharge of Ube2e2 was rapidly induced by a control E3 enzyme (RNF12^40^), but the DDD(ΔBPB) complex did not substantially increase the rate of ubiquitin discharge from Ube2e2 enzyme compared to the conjugate alone over an hour (Figure 2e). This suggests that binding to the DET1 cup in of itself does not destabilise the E2-ubiquitin conjugate, but associated E3s or other binding partners may promote an uncharged E2 in this position.

Given that the equivalent plant E2 protein that binds to DET1 (COP10) is catalytically inactive and cannot be charged, this raises the possibility that E2s binding to the DET1 cup may play a role other than ubiquitin transfer.

### Dynamic disorder of DET1

The process of 3D classification showed that DDD-complexes have variable degrees of order (Figure 3). Three-dimensional variability analysis and clustering revealed only a subset of particles that contained DET1 and DDB1 had density corresponding to the full DET1 protein (Supplementary Figure 2). Refinement of the most complete cluster led to the structure described in Figures 1 and 2. Another set of particles (Cluster 1; refined to 3.0 Å) had fully resolved DET1 but no Ube2e2 was visible, and there was no density corresponding to the cup of DET1. An even larger set of particles showed relatively clear density for the section of DET1 that sits above the BPC domain of DDB1 (residues 12–160, 449–550), but little density for the section that sits above the BPA domain (161–448). Further evidence for this dynamic behaviour was present in the data collected for the DDD complex with full-length DDB1, where data anisotropy prevented detailed model building, but DET1 in general appeared to be in a relatively open state (Supplementary Figure 1c). The difference between the clearly defined density of DET1 and the variable density of the portion between residues 161-448, strongly suggests that this region of DET1 is flexible, and may undergo dynamic unfolding to move between a ‘closed’ state and a more open state.

**Figure 3:**
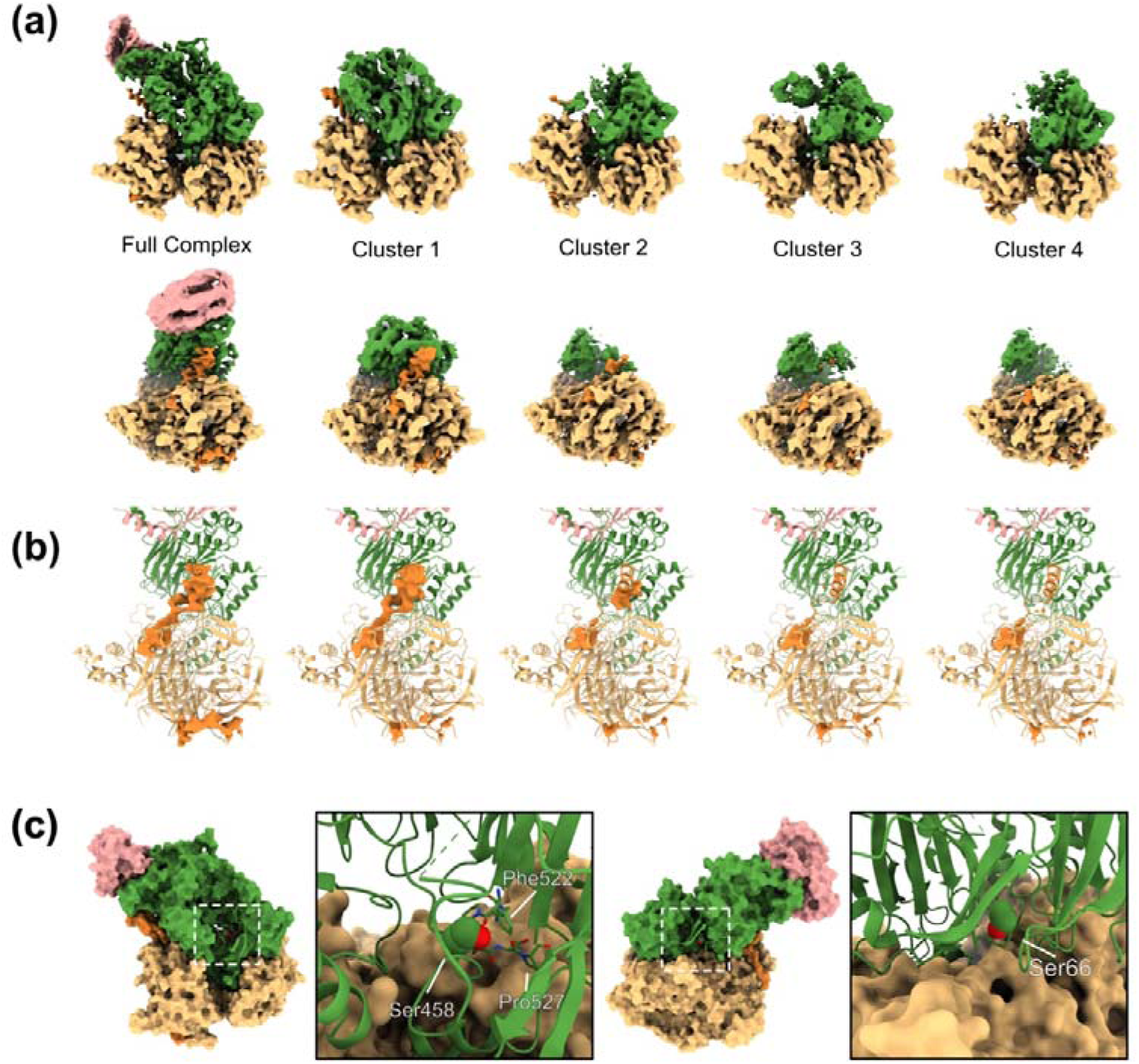
The DDD(ΔBPB) complex exists in a variety of discernible density states. 3D classification and 3D variability analysis of CryoEM data for the DDD(ΔBPB) complex reveals multiple subclasses that are indicative of a dynamic complex **(a)** Structures of variable DET1 order from different clusters (Supplementary Figure 2b). Complexes with density for all the components of the DDD(ΔBPB) complex define the “closed” state. As the complex shifts from a closed to an open state, density becomes more sparse. More complete models contain more density for DDA1. **(b)** Corresponding carved DDA1 density across different classes in *Panel a*. The full complex and cluster 1, the most complete models displayed, contain more density for DDA1 than clusters 2-4, where DDA1 density is disparate, indicating that DDA1 may lock the complex into the closed state. **(c)** Phosphorylation of DET1 may prevent a closed conformation. Ser66 and Ser458 of DET1 are subject to phosphorylation. Phosphorylation of Ser66 and Ser458 would preclude the structure from forming the closed state. Insets show the position of Ser458 and Ser66 as spheres.

In the structures where DET1 is closed (Figure 3; Full structure, Cluster 1) the C-terminal helix of DDA1 clearly interacts with DET1 (Figure 3). However, in the more open classes there is disparate density for DDA1 (Figure 3a/b). This could be explained by the fact that both DDA1 and DET1 are dynamic. However, when viewed at equivalent contour levels in the DDB1 region, the more open classes lack density corresponding to either N- or C-terminal portions of DDA1 (Figure 3b); this suggests that the most disordered classes have low overall DDA1 occupancy. Regardless of whether this is a cause or effect of DET1 or DDA1 dynamics or low occupancy, the order of DDA1 is closely linked to stability of the closed conformation of DET1.

Previous studies have reported the regulation of DET1-COP1 substrate recruitment by MAP kinases^10^. Zhang et al reported two prominent phosphorylation sites on DET1—Ser66 and Ser458—which modulate its ability to degrade substrates in co-operation with COP1. The flexibility of DET1 we observe provides an insight to how this could occur. Ser458 sits at the edge of the dynamic region of DET1. In the closed conformation this region folds back onto itself to interact with the core of the molecule. The other MAPK target residue, Ser66, is similarly located within the buried DET1 core. Un-phosphorylated Ser458 interacts with the loop containing Phe522 in the DET1 core. (Figure 3c).

Dynamic opening and closing of DET1 may allow Ser458 to be phosphorylated within the intact DDD complex however, Ser66 is likely only able to be phosphorylated when DET1 is dissociated from DDB1. The tightly packed arrangement of the closed conformation would not be compatible with a phosphorylated Ser66 or Ser458 in these locations (Figure 3c).

Thus, the central position of phosphorylatable residues within closed DDB1-DET1 suggests that they likely destabilise the closed conformation, and so may regulate DET1 closure.

### Closed DET1 underlies COP1 binding

As phosphorylation regulates DET1-COP1-mediated substrate degradation^10^, and potentially regulates the closed conformation of DET1; we hypothesised that DET1 closure and COP1-binding could be linked. To test this, we turned to AlphaFold2 to explain how the closed conformation of DET1 may impact DET1-COP1 co-operativity. We first modelled a 1:1 complex of DET1 with near full-length COP1, omitting the N-terminal 125 residues which are highly disordered in most predicted COP1 models. The highest scoring predictions showed COP1 binding to DET1 via its WD40 domain in two different modes (designated COP1’ and COP1’’), each with high confidence predicted aligned error (PAE) scores (<5 Å) for interface residues (Supplementary Figure 6). Both binding modes were largely compatible with each other, so we tested modelling DET1 bound with two copies of COP1. All 25 models of a 1:2 DET1:COP1 complex show a near-identical arrangement of the COP1 dimer bound to DET1, with correspondingly high iPTM scores and PAE of interface residues <5 Å (Figure 4a/b; Supplementary Figure 6). In these models, the COP1 WD40 domains occupy both of the binding sites identified in the 1:1 models, and the COP1 coiled-coil regions form an anti-parallel arrangement. The two WD40 domains of the COP1 dimer bind to distinct interfaces on DET1 through non-equivalent regions of their WD40 domains.

**Figure 4:**
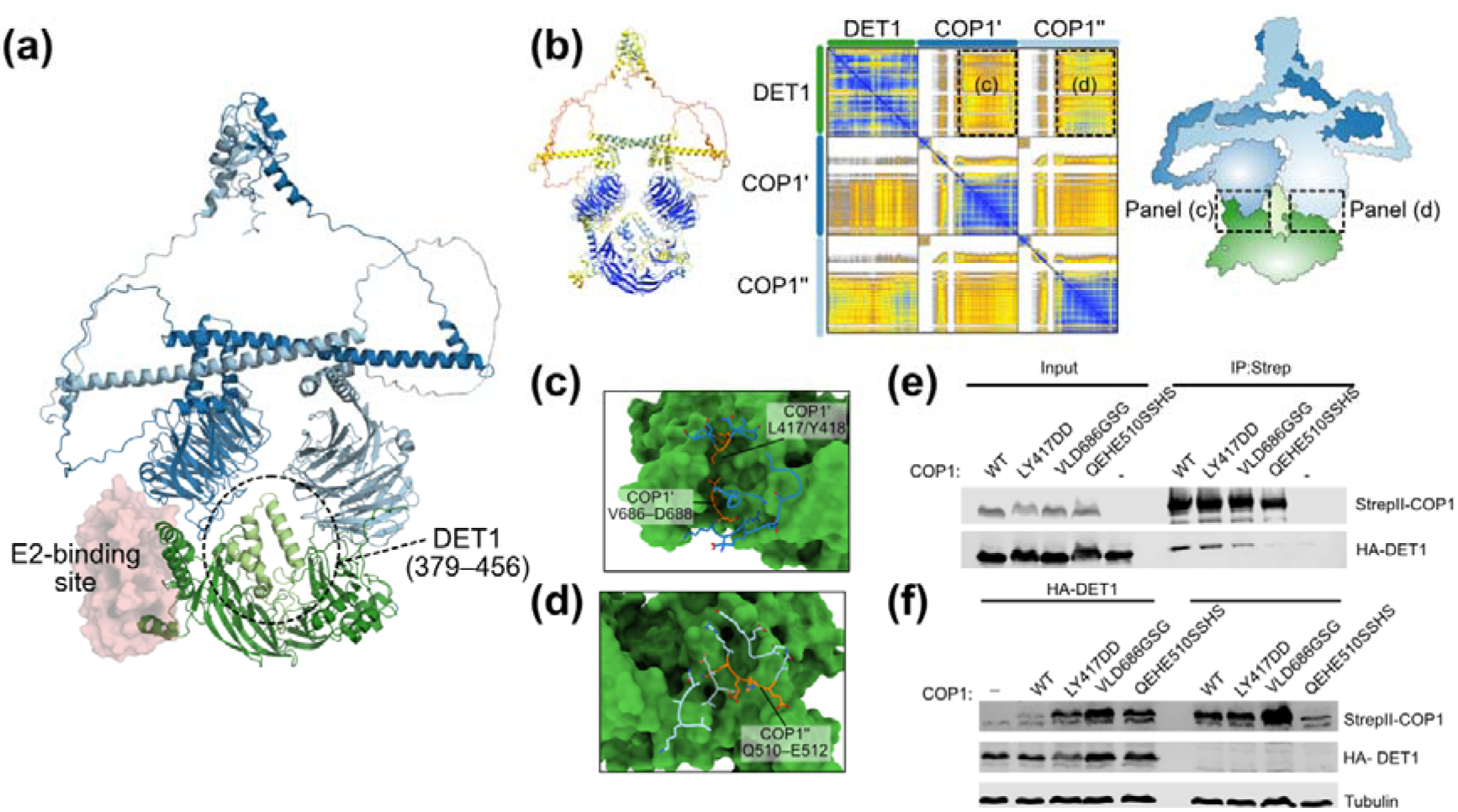
COP1 recruitment by DET1. **(a)** AlphaFold model of 2:1 COP1-DET1 complex, indicating dynamic region and E2-binding cup. **(b)** PAE plot, and overall LDDT scores relative to **(c/d)** Closeup views of the COP1’ and COP1’’ interfaces with DET1 **(e)** Mutations of COP1 targeted to disrupt interactions with DET1 show that COP1” is important for binding of COP1 to DET1. Western blot analysis was used to determine co-purification of HA-DET1 by either WT COP1, or binding mutants -COP1 LY417DD, VLD686GSG, (COP1’) or QEHE510SSHS (COP1”) following affinity purification with StrepTactin®XT 4Flow resin. **(f)** Co-expression of DET1 and COP1 in Expi293 cells shows destabilisation of wild-type COP1 by DET1 but not by COP1’ or COP1” mutants. Western blotting was performed to determine COP1 and HA-DET1 levels.

The first COP1 binding site on DET1 (COP1’) is adjacent to the Ube2e-binding site and is primarily mediated by two loops from the COP1 WD40 domain (Figure 4c). The second COP1 binding site (COP1’’) is nearer the N-terminal portion of DET1 and is mediated by multiple different loops within the WD40 domain (Figure 4d). The two DET1-binding sites are relatively comparable in area (1174 and 938 Å^2^ for COP1’ and COP1’’, respectively); however, both WD40-DET1 interfaces are substantially smaller than the area buried by COP1 self-association within the dimer (3242 Å^2^). While the precise details of the AlphaFold interfaces should be viewed conservatively, the overall characteristics are consistent with a COP1 dimer binding to DET1 through two asymmetric binding sites of DET1.

Both the COP1 binding-interfaces of DET1 are centred on a portion of DET1 (379-456) that is stabilised in the closed conformation, but not in the more open subclasses of DET1 (Figure 3). To test the AlphaFold2-based model for DET1-COP1 binding we designed COP1 mutations that would affect each interface, expressed COP1 and DET1 separately, and then tested the ability of the mutants to co-purify with DET1 (Figure 4e). Within COP1’ we mutated Leu417/Tyr418 and Val686/Leu687/Asp688 and within COP1’’ we mutated Gln510/Glu511/His512/Glu513. Mutations of COP1” markedly reduced the amount of DET1 which could be co-purified by COP1. In contrast, mutations in COP1’ had a more moderate effect on binding. The Leu417/Tyr418 mutant that predominantly faces the E2-binding site showed roughly wild-type binding, while the Val686/Leu687/Asp688 mutant had reduced binding relative to WT COP1 (Figure 4c).

To further validate the complex we co-transfected Expi293 cells with either wild-type COP1 or each of the loop mutants and/or full-length DET1. Co-transfection of wild-type COP1 and wild-type DET1 resulted in a significant destabilisation of COP1, relative to when wild-type COP1 was transfected alone, presumably because COP1 is being recruited to the CRL4^DET1^ complex and being proteasomally degraded. While we cannot verify that the primary role of DET1 is degradation of COP1 at endogenous levels in biologically relevant cell lines, this effect does serve as a useful readout for formation of CRL4^DET1-COP1^ complexes. When we co-transfected COP1 mutants at either COP1’ or COP1’’ interfaces COP1 levels were comparable to the levels of each protein without DET1 present, indicating they are no longer are recruited to CRL4^DET1-COP1^ (Figure 4f). Overall, binding data suggests that the COP1’’ interface may be most crucial for initial DET1 binding, but mutation of either COP1’ or COP1” disrupt incorporation into CRL4^DET1-COP1^.

### Ube2e2 controls CRL4^DET1–COP1^ complex assembly and COP1 stability

With a strong structural model of assembly of the CRL4^DET1-COP1^ complex, we aimed to understand the role of Ube2e2 in the complex. To interrogate the effect on COP1 of removing Ube2e enzymes from CRL4^DET1^ we designed a DET1 ‘cup deletion’ mutant, that completely deletes the residues (264-327) required for E2 recruitment. We expressed DET1 (either wild-type or cup deletion mutant) and COP1 separately, mixed the lysates and assessed the amount of DET1 which could be co-purified by COP1. Retention of the DET1 cup-deletion by COP1 was significantly reduced as compared with that of the full-length DET1, suggesting that the Ube2e-binding cup promotes assembly of CRL4^DET1-COP1^ complexes (Figure 5a). To test the consequences of E2 incorporation we co-transfected Expi293 cells with WT (full-length) COP1 and either full-length DET1, or the cup-deletion mutant. In the presence of cellular CUL4 and ubiquitin ligase machinery, co-transfection of wild-type DET1 and COP1 saw a significant destabilisation of COP1, relative to when COP1 was transfected alone. This suggests that COP1 is being recruited to the CRL4^DET1^ complex but being ubiquitinated by CRL4^DET1^ and destabilised. In contrast, when COP1 was transfected with the DET1 cup-deletion mutant the COP1 is present at near the same level as the COP1 which was transfected alone (Figure 5b). This suggests that the residues which are important for DET1 to bind to Ube2e2, and so E2 recruitment, is also important for destabilisation of wild-type COP1 in Expi293 cells.

**Figure 5:**
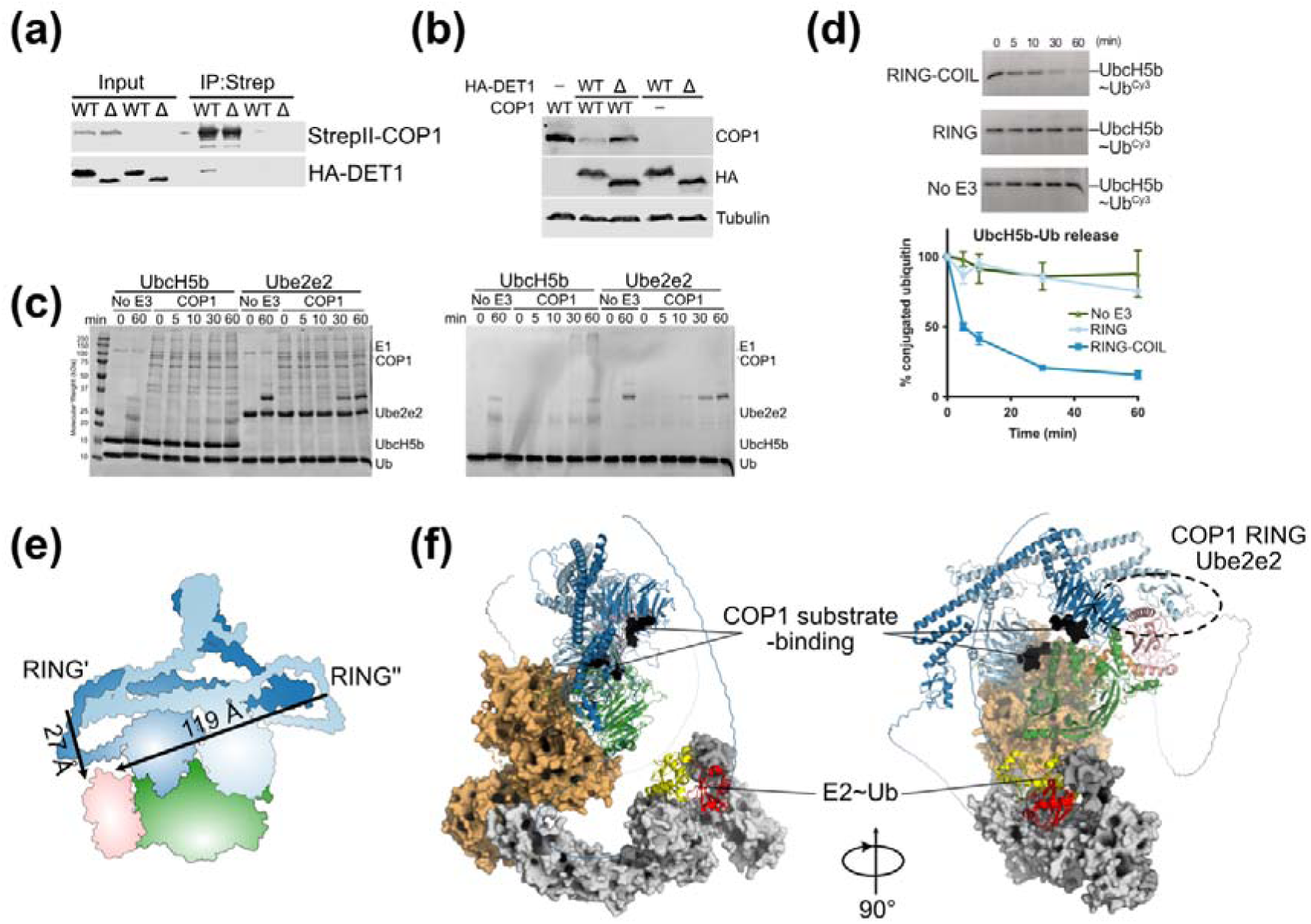
Activity and regulation of COP1 activity. **(a)** Pulldown of DET1 and DET1 cup deletion mutant (Δ) by COP1. COP1 interacts with full-length DET1 but not the DET1 which lacks the cup. Co-purification of HA-DET1 (either WT or Δ) by StrepII-WT COP1 following Strep affinity purification was assessed by Western blotting. **(b)** Wild-type COP1 is destabilised by full-length DET1 but not the DET1 cup mutant. Expi293 cells were co-transfected with StrepII-COP1 and either HA-DET1 or the DET1 cup mutant. Western blotting was performed to determine COP1 and HA-DET1 levels. **(c)** COP1 is active with UbcH5b but not Ube2e2. Multi-turnover activity assays were used to determine the ability of COP1 to build ubiquitin chains with either UbcH5b or Ube2e2. Samples were taken at 0, 5, 10, 30 and 60 mins and ubiquitin chain formation determined by visualising the incorporation of Cy3-labelled ubiquitin (right) and the gels stained with Coomassie Blue (left). **(d)** Ubiquitin discharge assays were used to determine the ability of COP1 RING or COP1 RING-COIL to cause ubiquitin discharge from UbcH5b. Samples were taken at 0, 15, 30 and 60 mins and ubiquitin discharge was determined by visualising Cy3-Ubiquitin (e) Location of Ube2e2 in the DDD(ΔBPB) complex position relative to COP1 RING domains. The arrangement of COP1 in an antiparallel dimer and the DDD(ΔBPB) complex would preclude a COP1 RING dimer. **(f)** Hybrid model of the CRL4^DET1^-COP1-Ube2e2 complex. Based on superposition of an Alphafold3 model of DDD-COP1-Ube2e2 a model was created by sequential superposition of the CUL4-DDB1-DCAF1 structure, and the active CUL1 structure, and COP1-domain bound to TRIB1 peptide. The low-LDDT tail of Ube2e2 is omitted for clarity, E2∼Ub shown in yellow-red, and substrate binding site of COP1 in black.

To further understand if Ube2e enzymes may mediate COP1 destabilisation we tested basal activity of full-length COP1 with Ube2e2 and UbcH5b, a promiscuous E2 protein. Multi-turnover ubiquitination assays in the absence of the DDD complex showed that COP1 was highly active in autoubiquitination with UbcH5b, but inactive with Ube2e2 (Figure 5c).

Similar results were seen for COP1 activity on model substrates, with no measurable activity with Ube2e2 (Supplementary Figure 7). These results show that full-length COP1 alone does not display activity with Ube2e2, even though it is active with a promiscuous E2. Similar lack of activity with Ube2e2 was observed when DDB1-DDA1-DET1 were also included in COP1 ubiquitination assays (Supplementary Figure 8). Therefore, the role of Ube2e2 in the DDD complex is unlikely to be promoting COP1 autoubiquitination but may have a different role entirely.

An alternative role for Ube2e within CRL4^DET1-COP1^ complexes could be to stabilise COP1 within the complex, but suppress COP1 RING domain activity. Because RING domains frequently require dimerization for activity, we tested the impact of COP1 oligomeric state on COP1 RING activity. We used multiple angle-light scattering and observed that the COP1 RING domain (residues 126–208) in isolation forms weak, concentration-dependent dimers (Supplementary Figure 9). The RING-only construct also had minimal activity in E2-ubiquitin discharge assays, whereas a COP1 RING-coil (residues 126–308) construct had significantly higher activity. This would be consistent with the helix C-terminal to the RING mediating dimerization and promoting activity (Figure 5c). However, an anti-parallel arrangement of the COP1 coiled-coils suggested by Alphafold would mean that the RING domains must span the coiled-coils to be active within a dimer. In the CRL4^DET1-COP1^ model, one COP1-RING domain would be adjacent to the RING binding surface of Ube2e2— approximately 27 Å from the end of the coil to the E2-binding site, which could be comfortably bridged by the RING-COIL linker. However, the other RING domain would need to span approximately 120 Å from the predicted end of the coil (Figure 5d). This means that the role of Ube2e2 in the CRL4^DET1-COP1^ may be to sequester a single COP1 RING domain, preventing RING dimerization rendering it less active. Such a mechanism would allow the activity of CRL4 to predominate and mediate ubiquitination of substrates or COP1 itself, depending on the biological context.

### Model of the CRL4^DET1^-COP1 holo-complex

While this manuscript was in its final stages of preparation, Alphafold3 enabled modelling of the entire CRL4^DET1^-COP1-Ube2e2 complex^41^. Alphafold3 predictions are extremely consistent with the proposed model—it reiterates that a closed conformation of DET1 is required for COP1 binding, and the interfaces between DET1, Ube2e2, and COP1 are near identical to those identified in cryo-EM data, predicted by Alphafold2, and validated by experimental data. In ten replicate runs the COP1 RING domain occupies the canonical RING-binding site of Ube2e2, although PAE scores are relatively moderate (Supplementary Figure 10). Alphafold3 does not provide any confident predictions about the relative positions of COP1 and CUL4 however, so we created a hybrid model combining solved structures of NEDD8-modified CUL1 in the act of transferring ubiquitin^42^, and full length CUL4 bound to DCAF1^19^, and the COP1 WD40 domain bound to COP1^31^. This clearly illustrates DET1 closure and subsequent binding of COP1 would orient CRL4^DET1^-COP1 well for activity. Closed DET1 orients both substrate-binding pockets of COP1 directly towards the Rbx1 RING domain, upon which a charged active ubiquitin-conjugating enzyme would be poised to execute ubiquitination (Figure 5f).

## DISCUSSION

Amongst the myriad of eukaryotic ubiquitin E3-ligases, Cullin-RING ligases recruit diverse substrates by building a modular system upon a common core. DCAF proteins provide diversity of Cullin-4 substrate recruitment when they are recruited via DDB1 and are particularly relevant to transcriptional regulation and DNA-damage responses. Of the DCAF proteins, DET1 is somewhat enigmatic, given its primary reported functions have been inhibition of CRL4 ubiquitination, binding to E2 ubiquitin-conjugating enzymes^28^, and recruitment of another E3 ligase protein to CRL4 complexes, namely COP1. Here we describe the molecular architecture of DDB1-DDA1-DET1-E2 complexes and characterise a model for recruitment of COP1 to CRL4^DET1^ complexes.

One of the most intriguing aspects of DET1 is its ability to recruit additional E2 enzymes to CRL4 complexes. This function appears to be unique amongst DCAF proteins reported to date. Published data and our analysis suggests that specific recruitment of Ube2e enzymes is mediated by a helical hairpin extending from DET1 that binds the backside beta-sheet of Ube2e2. Contacting residues of Ube2e2 are highly conserved with Ube2e1 and Ube2e3, and modelling suggests that this binding mode is crucial to specific recruitment. The binding site for DET1 on Ube2e2 clearly precludes backside-binding of ubiquitin^36,39^, which activates ubiquitin transfer of other E2 enzymes. We did not observe enhancement of ubiquitin discharge from Ube2e2 by DET1 in up to an hour; suggesting that backside binding alone is not activating. This unique DET1-Ube2e partnership adds to a number of Ube2e peculiarities in the literature. Ube2e enzymes preferentially catalyse mono-ubiquitination over poly-ubiquitination even though they avidly bind to BRCA1-BARD1^43^. The Ube2e N-terminal extension has been shown to impede extended ubiquitin chain building, and acts as a target of auto-ubiquitination that limits activity^44–46^. Moreover, the plant homolog of DET1 recruits a catalytically dead E2 enzyme (COP10), containing a serine rather than cysteine in its active site^29,30^. A catalytically dead E2 in COP10 would effectively enforce association of uncharged Ube2e1/2/3 with DET1 complexes, as observed in humans. These observations, combined with our data, contribute to a model where Ube2e family E2s are not primarily required for building of ubiquitin chains by COP1, but rather act to recruit COP1 to DDD complexes.

The analysis of dynamic regions of DET1 by cryoEM provides insight into potential regulation by phosphorylation. However, it also extends the paradigm of Cullin substrate adaptors proteins being more than static substrate-recruitment modules, but dynamic and active players in ubiquitination. Studies have shown that the substrate adaptor protein Cereblon, a target of CELmod compounds and a range of proteolysis targeting chimera molecules, exists across a spectrum of conformations. These conformations can be biased by different small molecules^33^. Furthermore, DCAF1 mediates assembly of massive tetrameric CRL4 complexes, which have low activity but become active when substrate-binding promotes dissociation into dimeric structures^19^. In the case of DET1, the fully-closed state generates two asymmetric binding sites that we propose mediate COP1 binding, in concert with the presence of Ube2e enzymes. The fully-closed structure of DET1 also correlates with stabilisation of the C-terminal helix of DDA1. DDA1 resides in a similar location to that observed in multiple structures reported of DCAF15 bound to DDB1^34,35^. While this has not been systematically reported, this may mean that DDA1 is also required for stabilisation of substrate bound DCAF15. Beyond DCAF15, the only other DCAF-DDB1 structures incorporating DDA1 is that of DCAF16^47,48^. In those structures, only the N-terminal portion of DDA1 is visible bound to the BPC WD40 domain, if at all. Moreover, DCAF16 appears poorly disposed to interact with DDA1. It will be important in the future to systematically ascertain which DCAF complexes DDA1 is an integral component of, and how DDA1 may modulate the function of respective DCAF conformations and compositions.

The model we present here for COP1 recruitment to CRL4^DET1^ sees a dimeric COP1 complex recruited to a monomeric DET1 protein through asymmetric binding sites. While AlphaFold2 also confidently allows for 1:1 complexes, the extended COP1 coiled-coil interface, that COP1 RING dimers are more active, and both 1:1 binding modes being compatible within a 2:1 complex all argue in favour of a 2:1 complex as the default oligomeric state. Within this complex, it appears unlikely that COP1 RING dimers would be favoured to come together into their active state, especially if one of the RING domains is bound to an Ube2e enzyme. This suggests that within the larger CRL4^DET1-COP1^ complex that COP1 activity is suppressed, and its role is restricted to that of a substrate recruitment module. A dimeric COP1 molecule binding to DET1 would present two WD40 domains with binding sites for substrates, or further substrate adaptors such as the Tribbles pseudokinases^7,31,49,50^. Reported COP1 substrates include multiple dimeric transcription factors (for example P53, C/EBP family, and Jun transcription factors). A dimeric ligase recruiting dimeric substrates presents an attractive matching stoichiometry. The role of Tribbles proteins as substrate adaptors within such complexes could offer a mechanism to introduce further diversity.

The question of why COP1 needs to be recruited to DET1 at all is still open, when COP1 can itself build ubiquitin chains on substrates. Despite the important roles of the COP1-DET1 targets and its role differentiation and development of multiple cell types, targeted proteomic studies have shown that DET1 and COP1 are amongst the least prevalent DCAF proteins resident within Cullin-4-DDB1 complexes^51^. This may mean that CRL4^DET1-COP1^ complexes acts to tune wider functions of COP1, or to extend chains on substrates more rapidly than COP1 is capable of itself. CRL4^DET1-COP1^ complexes may offer a route to control the rate of auto-vs substrate ubiquitination of COP1 or elevate rates of ubiquitination over COP1-only activity. Alternatively, CRL4^DET1-COP1^ may favour a particular subset of COP1 substrates over COP1 alone, or to control levels of substrate adaptors. The interplay between these different mechanisms represents the next step in understanding this key regulatory complex for developmental transcription factors.

## ACKNOWLEDGEMENTS

We acknowledge the financial support provided by the Marsden Fund (Grant UOO2102) administered by the Royal Society of New Zealand, which enabled us to pursue our investigations. TL and JD are supported by postgraduate PhD Scholarships awarded by the University of Otago, and LT is supported by a Postdoctoral Career Development Fellowship awarded by the Division of Health Sciences, University of Otago. Electron microscopy access was supported by a Royal Society of New Zealand Catalyst Seeding Grant (21-UOO-003-CSG). We also thank and acknowledge the use of the University of Wollongong Cryogenic Electron Microscopy Facility at Molecular Horizons. Special thanks to the Day Lab (University of Otago) for their collaboration and for generously sharing various DNA constructs and ubiquitin-related reagents, and Adam Middleton and Catherine Day for helpful comments on the manuscript. We also thank Peter Higbee (Research IT, Otago) for his expertise assistance which significantly contributed to the success of our project. We express our appreciation to the Electron Microscopy (EM) facility at Otago for their technical assistance.

## Supplementary Figures

**Supplementary Figure 1:**
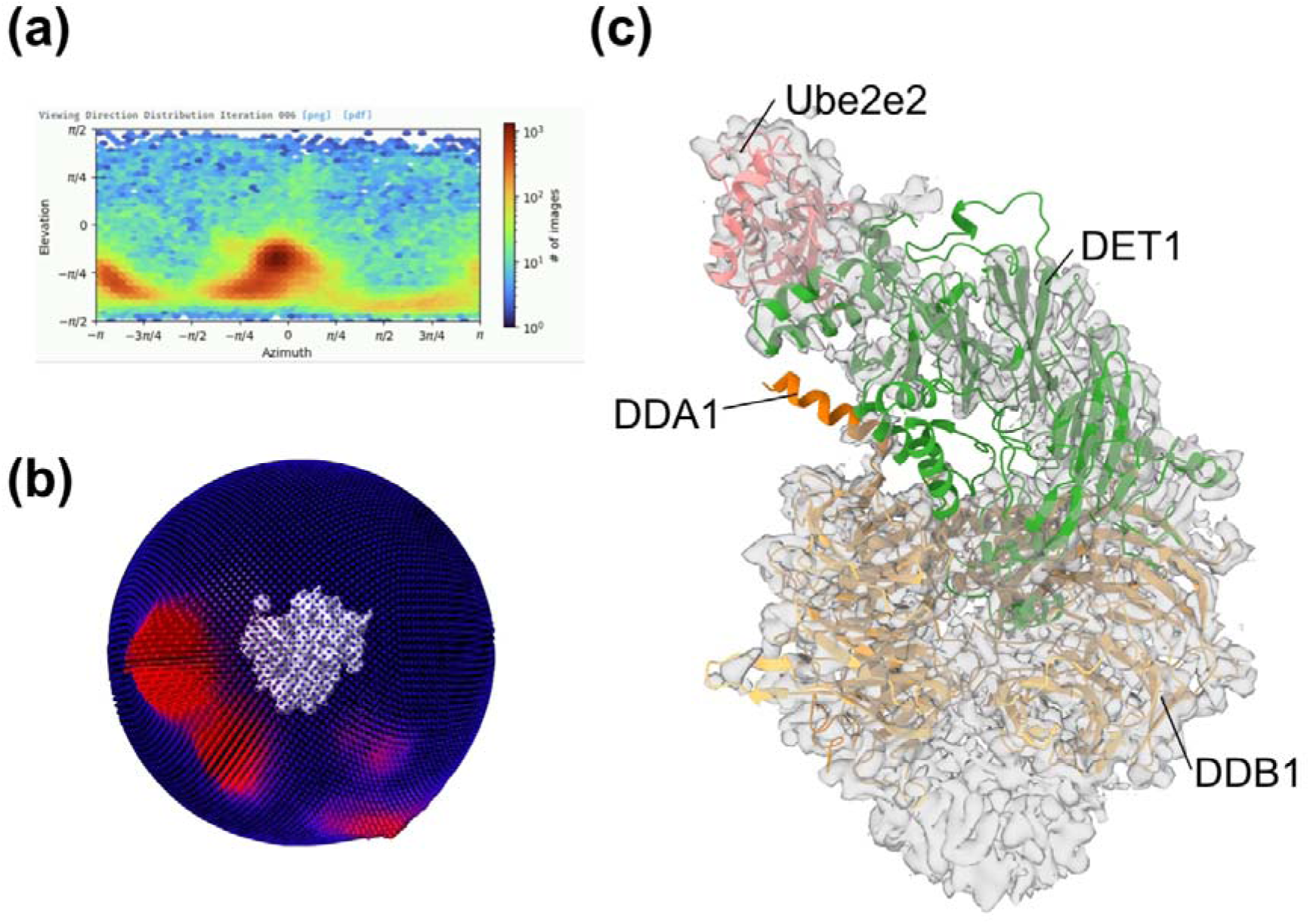
Initial structure of the full-length DDD:Ube2e2 complex showed anisotropy. **(a)** Direction distribution map shows severe anisotropy **(b)** Euler angle distribution **(c)** Cryo-EM density and molecular model of the DDB1(FL)-DET1-DDA1-Ube2e2 complex.

**Supplementary Figure 2:**
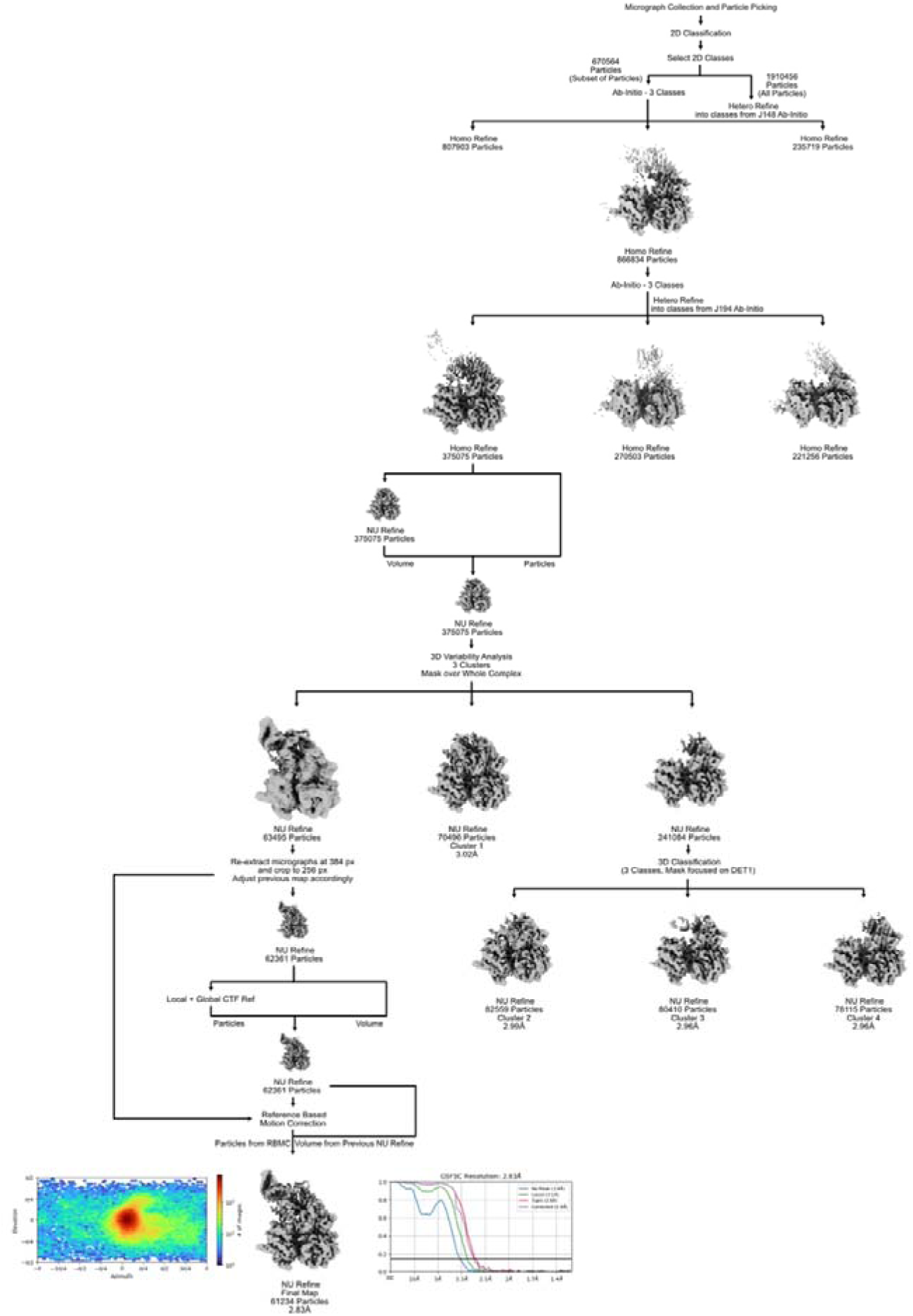
Cryo-EM image analysis workflow. Processing workflow described for the DDB1(ΔBPB)-DET1-DDA1-Ube2e2 complex. Motion correction, CTF estimation, particle picking, and particle extraction were all performed using cryoSPARC Live, before refinement in cryoSPARC. Initial 3D maps were generated *ab initio* from a subset of particles, before multiple steps of 3D classification, non-uniform refinement and 3D variability analysis were performed. Following 3D variability analysis, the micrographs of the best cluster were re-extracted at a higher resolution, before undergoing local and global CTF refinement. Finally, reference-based motion correction was performed and used in the final non-uniform refinement to generate the final reconstruction. Final reconstruction is shown at the bottom between angular distribution plot and 3DFSC. Clusters shown in Figure 3 labelled with resolution provided.

**Supplementary Figure 3:**
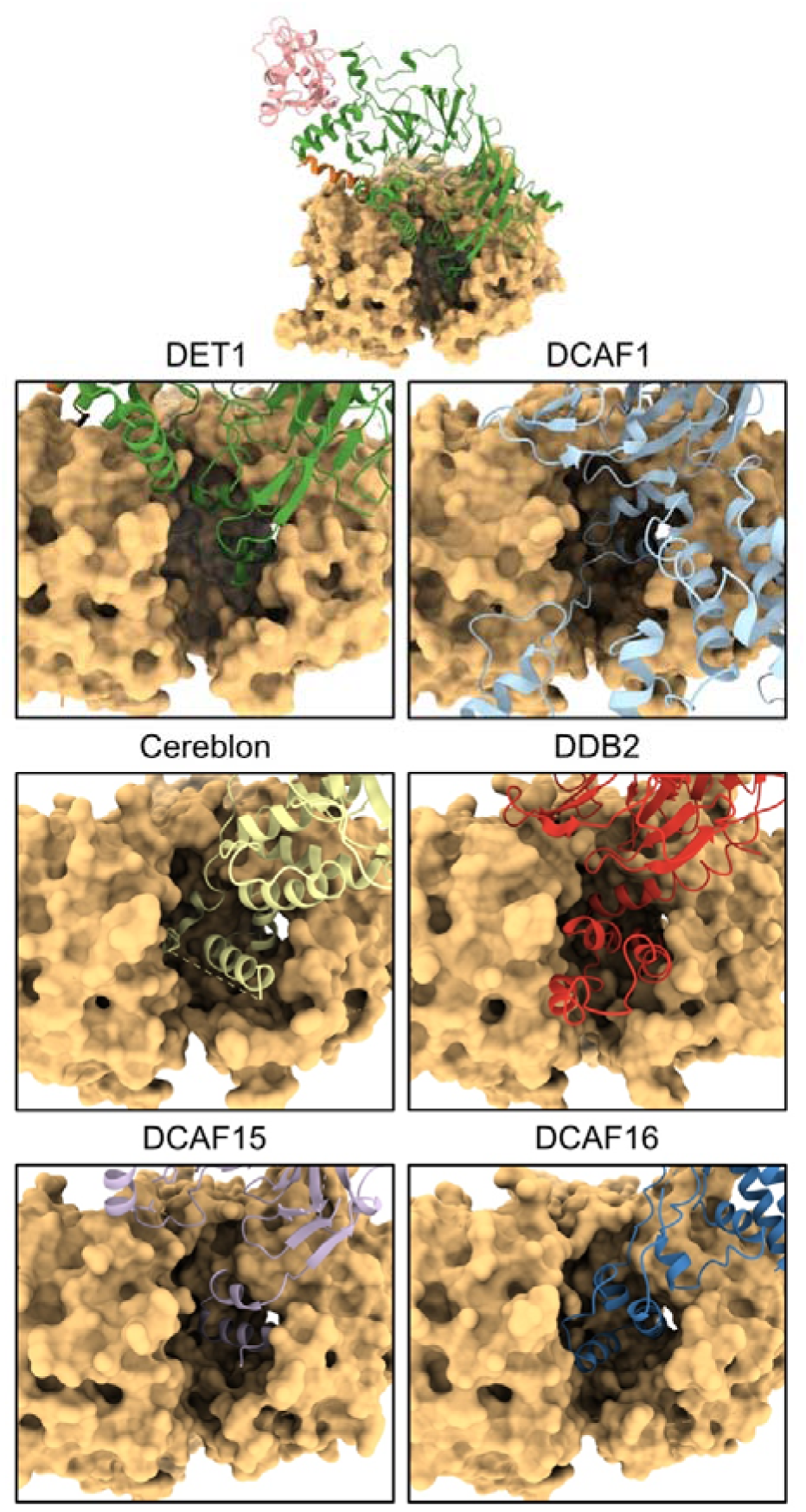
Comparison of DET1 and other DCAFs bound to DDB1. Previously solved structures of DCAFs binding to DDB1 were overlaid onto the cryoEM structure of the DDD (ΔBPB), displayed in Figure 1. All DCAFs bind to the same pocket of DDB1 as we observed for DET1. DCAF1 (PDB: 7V7C), Cereblon (PDB: 6BN7), DDB2 (PDB: 4E54), DCAF15 (PDB: 6PAI), DCAF16 (PDB: 8G46).

**Supplementary Figure 4:**
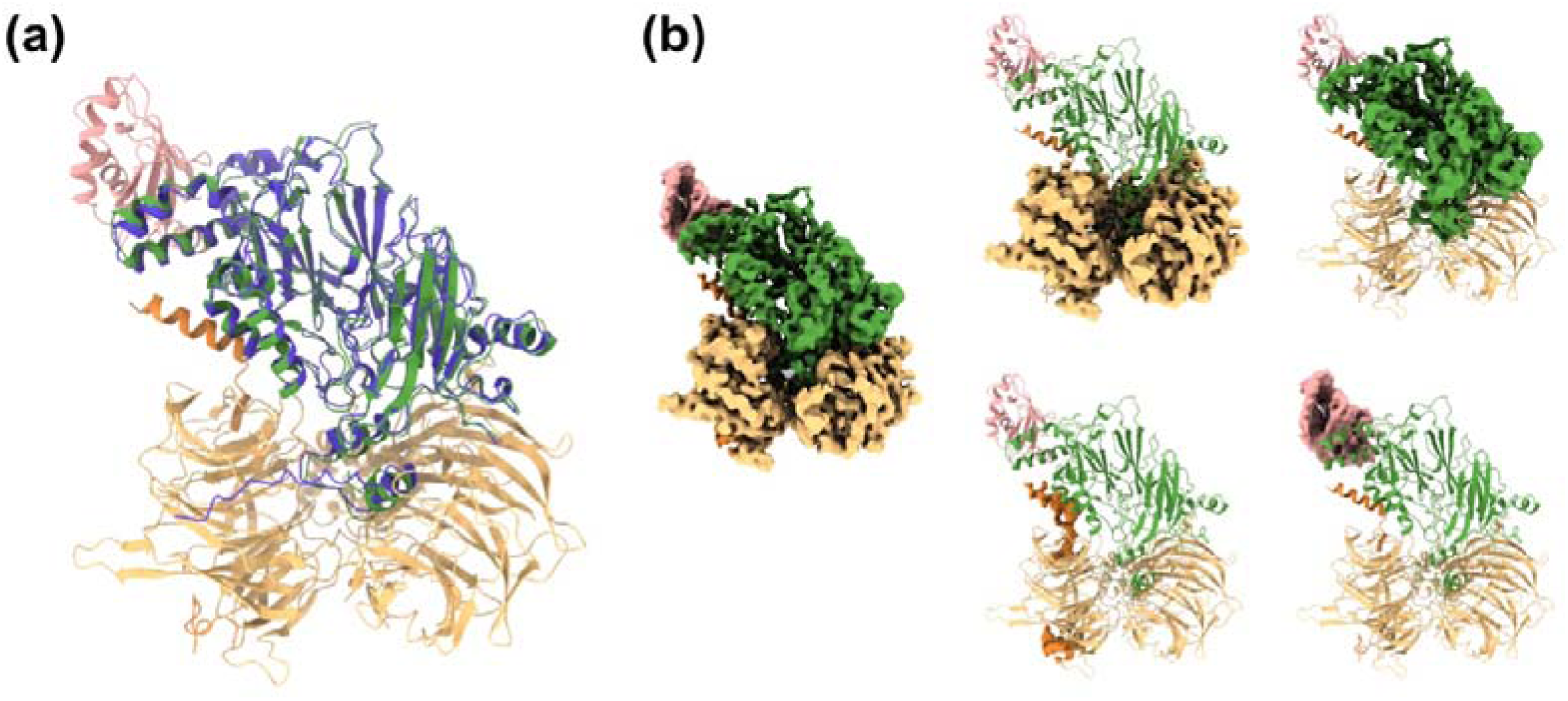
CryoEM structure of the DDD components. (a) Comparison of the AlphaFold model of DET1 to the cryoEM determined structure. The AlphaFold model of DET1 was overlaid onto the cryoEM structure of the DDD (ΔBPB), displayed in Figure 1 (RMSD of pruned atom pairs -0.931 Å, across all 508 atom pairs – 1.288 Å). DET1 AlphaFold model is displayed in purple. (b) Carved density for each component of the DDD (ΔBPB)-E2 complex. Density within 2.5 Å of the model was included in each component, with no overlap between components included; DET1 (green), DDB1(ΔBPB) (yellow), DDA1 (orange), Ube2e2 (pink).

**Supplementary Figure 5:**
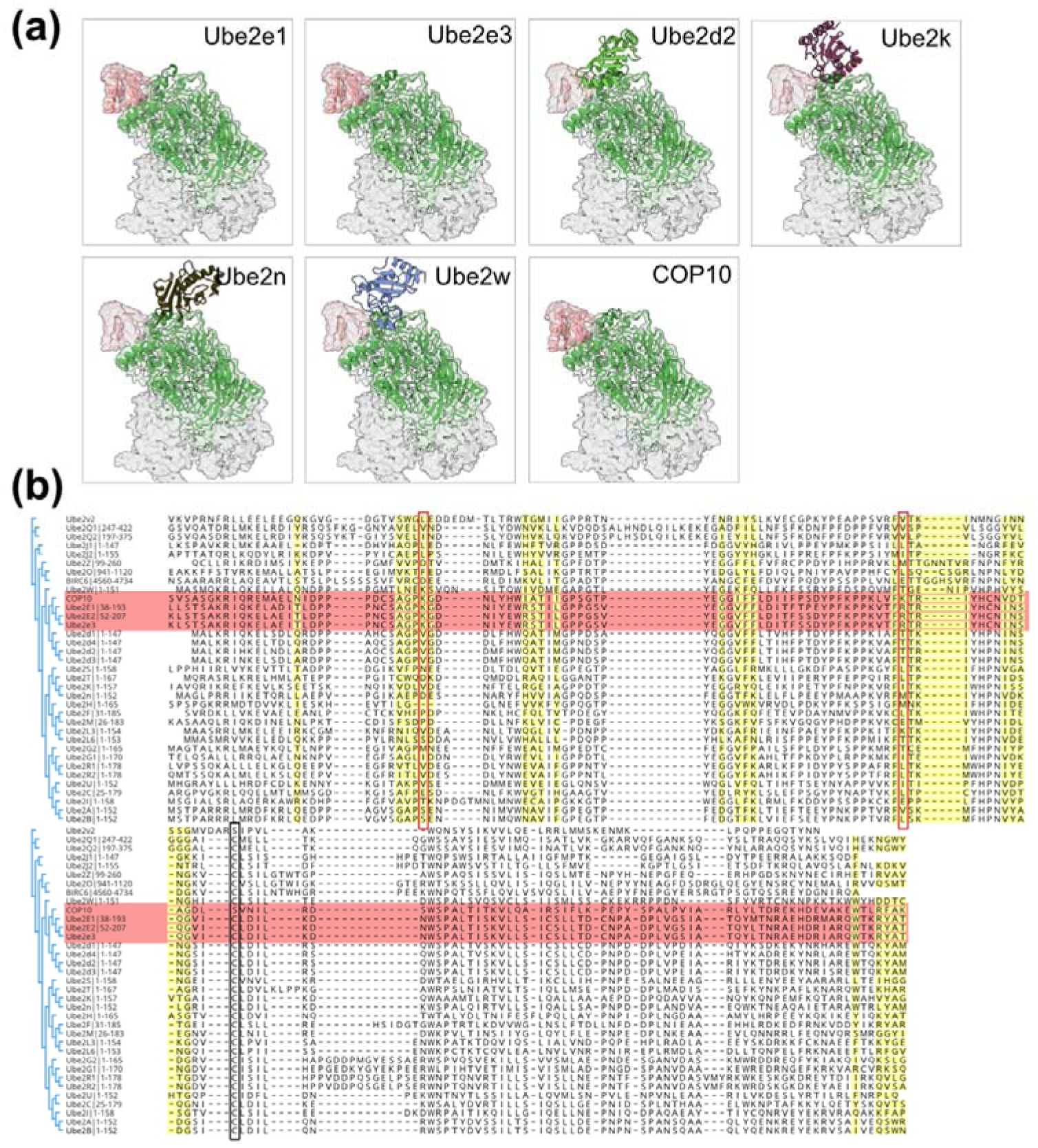
DET1 binds the Ube2e Family and COP10 using conserved residues within the E2 family. **(a)** AlphaFold models of various E2 enzymes show that only the closely related E2 enzymes, Ube2e1/3 and COP10, are compatible with the structure of DDD containing Ube2e2 (Figure 1) (b) Multiple sequence alignment of E2 enzymes with interface residues with DET1 highlighted in yellow. Red boxes indicate E2-DET1 interface residues divergent in the Ube2e family and COP10. The active cysteine is indicated by a black box.

**Supplementary Figure 6:**
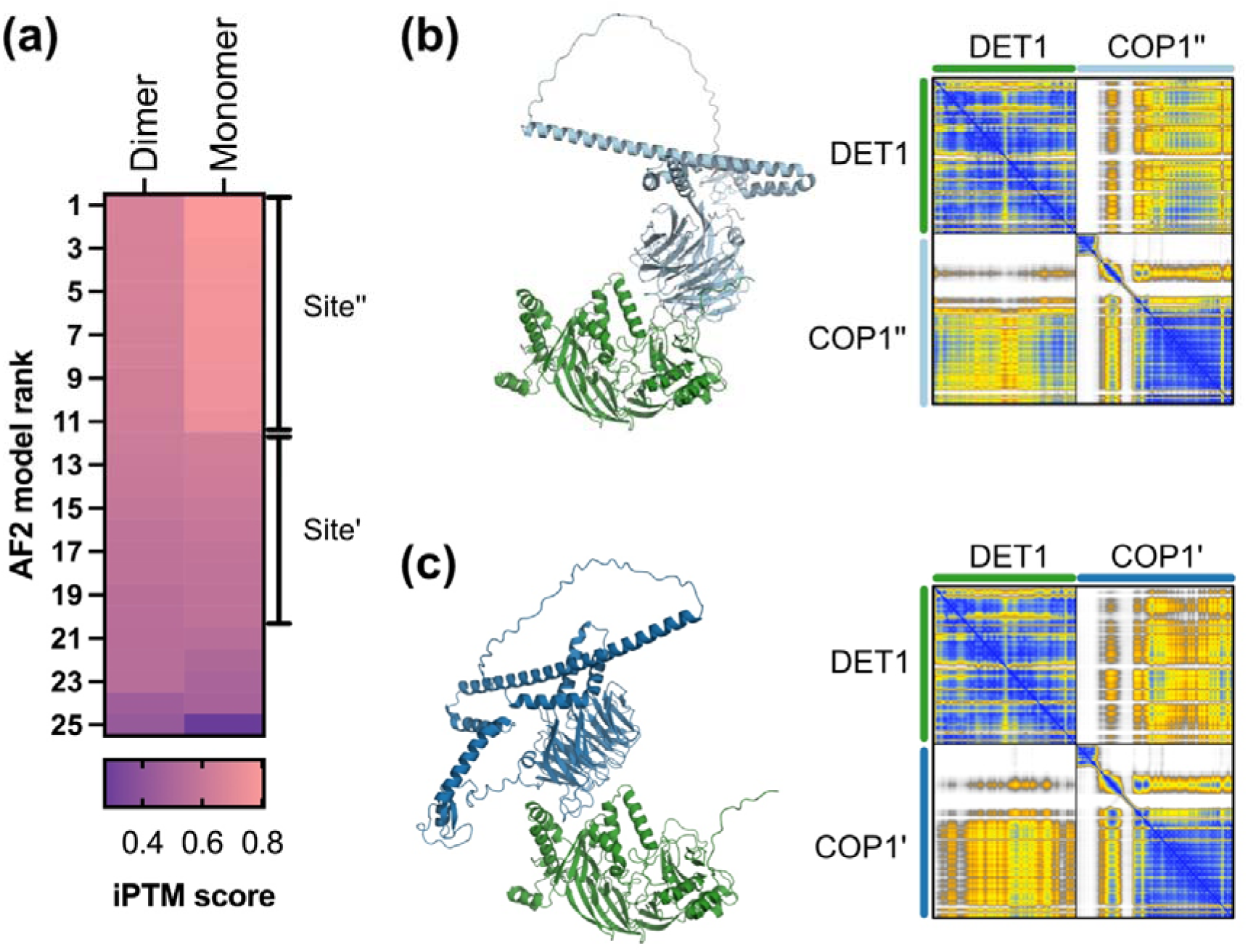
AlphaFold modelling of COP1-DET1 complexes. (a) AlphaFold models COP1-DET1 complexes with high confidence for both monomeric and dimeric COP1 binding to a single DET1. COP1 binds to DET1 through two distinct interfaces, (b) COP1’’ and (c) COP1’.

**Supplementary Figure 7:**
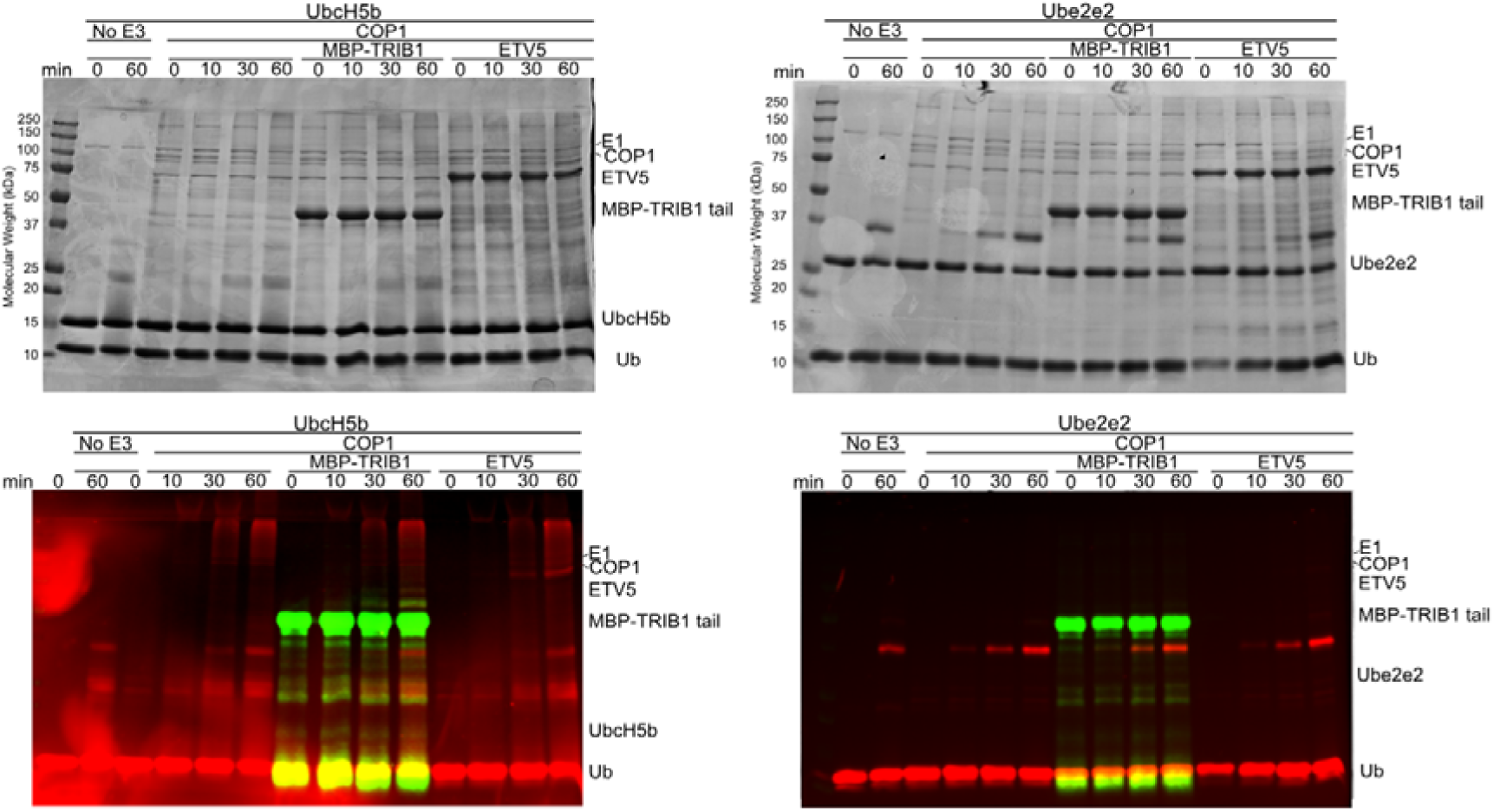
COP1 can act as an E3 with UbcH5b but not Ube2e2. Multi-turnover activity assays were used to determine the ability of COP1 to add ubiquitin onto substrates with either UbcH5b or Ube2e2. Samples were taken at 0, 10, 30 and 60 mins and ubiquitin chain formation determined by visualising the incorporation of Cy3-labelled ubiquitin (red) onto MBP-TRIB1^(345–372)^ (green, bottom) and the gels were stained with Coomassie Blue (top). UbcH5b allows for the incorporation of ubiquitin onto substrates which is demonstrated by the addition of ubiquitin on MBP-TRIB1 (green) and reduction of ETV5 (Coomassie).

**Supplementary Figure 8:**
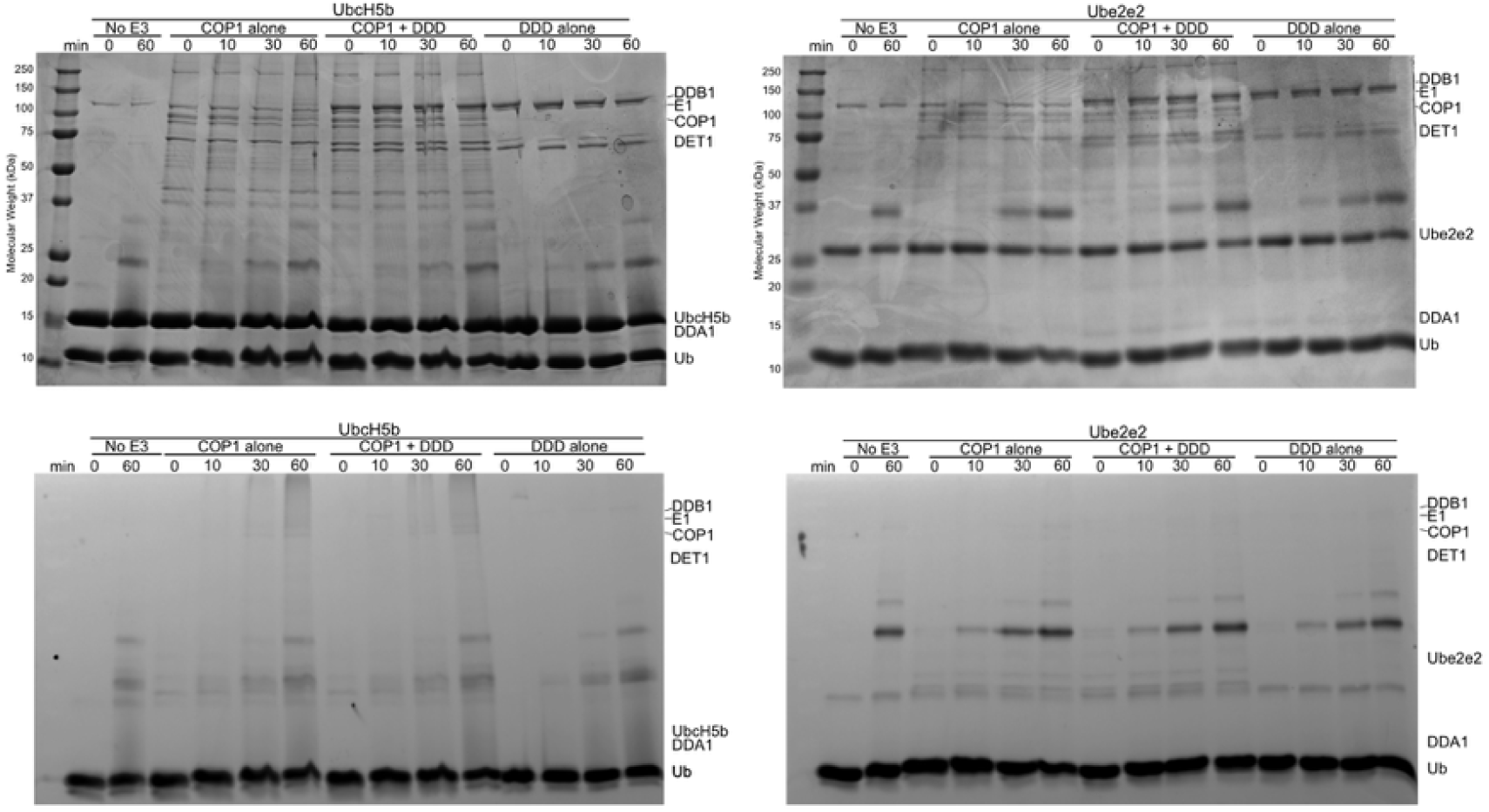
DDD does not affect COP1 activity. Multi-turnover activity assays were used to determine the ability of COP1 to add ubiquitin onto substrates with either UbcH5b or Ube2e2 in the presence of full-length DDD. Samples were taken at 0, 10, 30 and 60 mins and ubiquitin chain formation determined by visualising the incorporation of Cy3-labelled ubiquitin (bottom) and the gels were stained with Coomassie Blue (top). DDD has no intrinsic E3 activity alone, nor does addition of it affect COP1 activity with either UbcH5b or Ube2e2.

**Supplementary Figure 9:**
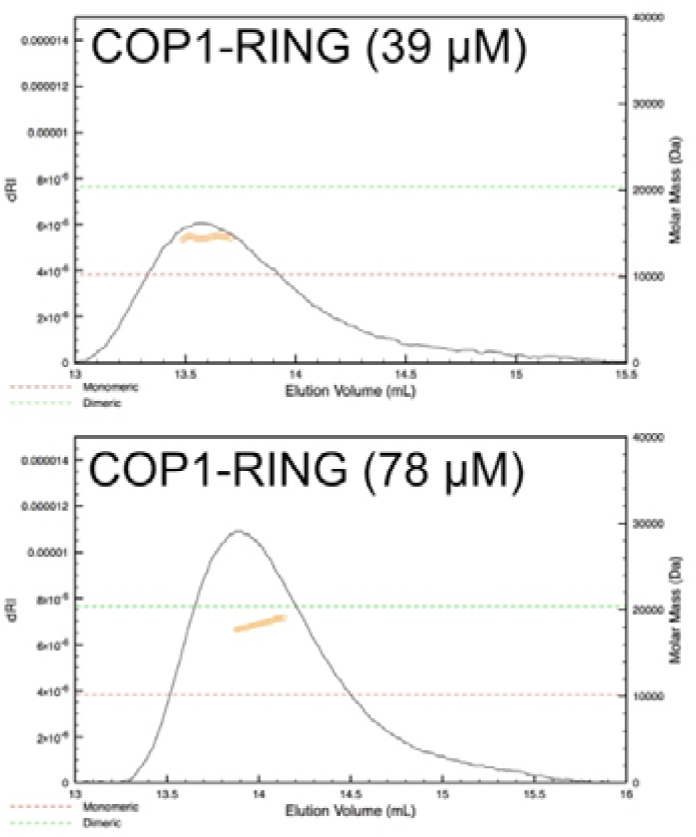
COP1 RING domain forms a weak dimer. Purified COP1 RING domain was analysed at a low concentration (39 µM) and high concentration (78 µM) by size exclusion chromatography multi-angle light scattering (SEC MALS). The theoretical molecular mass of a monomer (10.2 kDa, red) and homodimer (20.4 kDa, green) are indicated. The calculated molecular mass is indicated by yellow circles, either 14 kDa (at 39 µM) or 18 kDa (at 78 µM). The heterogeneity of the protein population is indicative of sampling between oligomeric states, which is concentration dependent.

**Supplementary Figure 10:**
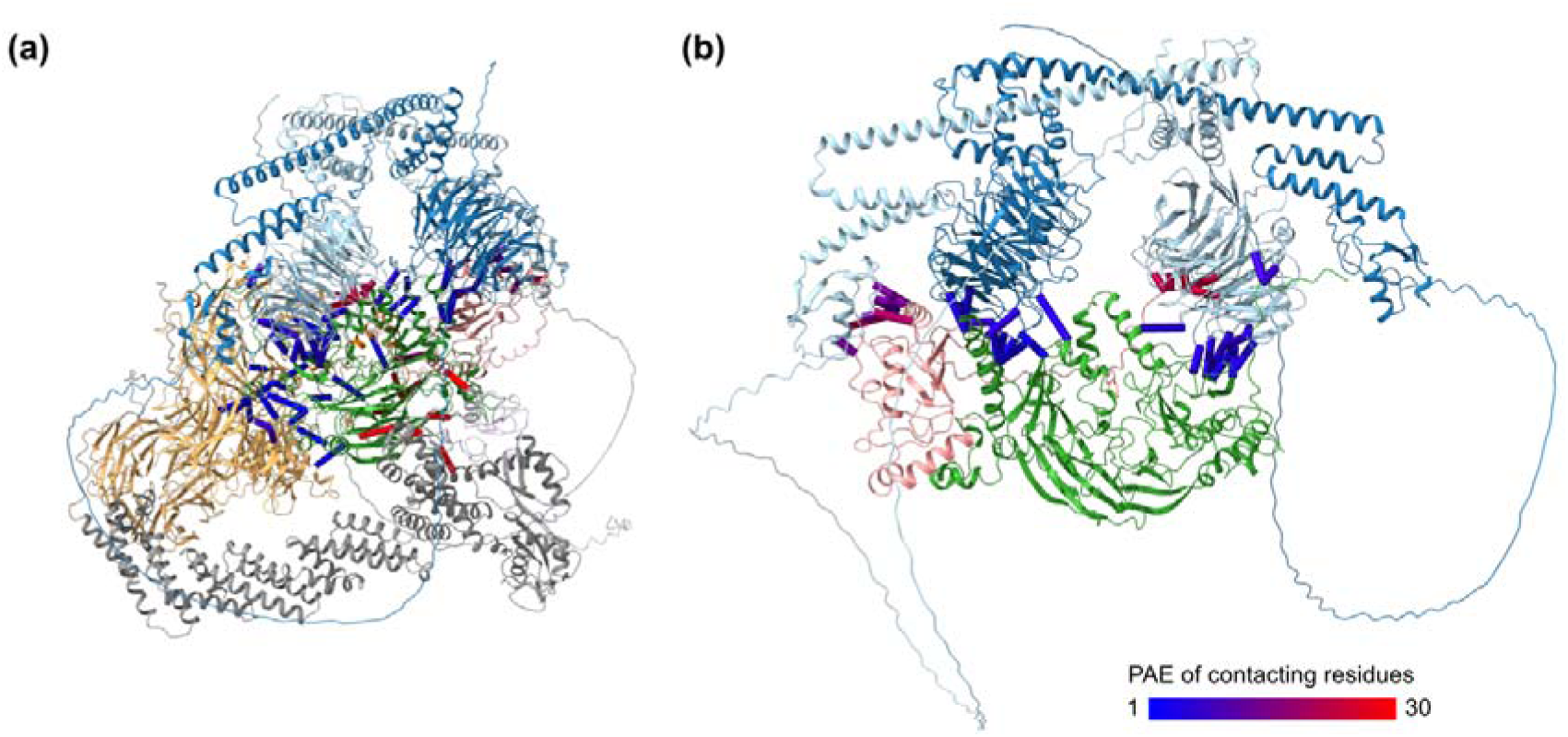
Alphafold3 model of the DDD complex. **(a)** Model of Cullin-4, Rbx1, DDB1, DDA1, DET1, a COP1 dimer and produced by Alphafold3. Contacting residues (within 3 Å of DET1 or Ube2e2) are shown with pseudobonds, and coloured according to PAE values. Ten runs with unique seeds produced effectively identical results. **(b)** Model from (a), with Cullin-4 and DDB1 omitted for clarity. Interfaces between COP1-DET1 are predicted with PAE <7 Å, and COP1 RING-Ube2e2 contacts with PAE 10–15 Å.

## MATERIALS AND METHODS

### Plasmids and cloning

DNA constructs for recombinant protein expression were amplified to incorporate the 5′- (CAGGGACCCGGT) and 3′-(TAACCGGGCTTCTCCTCG) overhangs, required for Ligation-Independent Cloning (LIC).

To generate a co-expression construct for the DDD complex, full-length constructs of DET1, DDB1 and DDA1were cloned into a pFastBac expression vector modified to incorporate an N-terminal affinity tag followed by a 3C[protease cleavage site. DDB1 and Ube2e2 were cloned to include an N-terminal His_6_ tag, DET1 an N-terminal StrepII tag and DDA1 remained untagged. These were then re-amplified and assembled into co-expression complexes in pBig1a using Gibson assembly. An additional construct was produced in which the BPB of DDB1 was removed (Δ396-706; ΔBPB). To then generate a co-expression bacmid construct, the pBig1a co-expression construct was transformed into DH10EMBacY cells (Geneva Biotech) and screened to identify constructs containing all the complex components. Full-length COP1 was cloned into the pFastBac expression vector modified to incorporate an N-terminal StrepII tag followed by a 3C[protease cleavage site and was transformed into DH10EMBacY cells to produce the bacmid. The isolated bacmids were transfected into *Spodoptera frugiperda* cells and used to produce baculoviral stocks which were then used for subsequent protein expression.

DNA constructs for protein expression in Expi293F (Thermo Fisher Scientific) were produced as follows: Full-length COP1was cloned into a modified pcDNA3.1 expression vector modified to incorporate an N-terminal StrepII tag and DET1 into the equivalent vector with an N-terminal HA tag. The DET1 cup mutant (ΔBPB) was generated by two-step overlap PCR and cloned into the same base vector as the full-length construct. Mutations in COP1were introduced by one-step site-directed mutagenesis.

COP1 RING and COP1 RING-COIL were cloned into a modified pET-LIC vector incorporating an N-terminal His_6_ tag and a 3C protease cleavage site for expression in *Escherichia coli* BL21 (DE3). Plasmids encoding the various proteins required for ubiquitination, including the Ube1 (E1), various E2s and Ubiquitin were provided by the Day Lab (University of Otago). ETV5_pet28a was a gift from Peter Hollenhorst (Addgene plasmid # 131639)^52^.

### Expression and Purification of Recombinant Protein

Bacterial constructs (COP1 RING/RING COIL, and proteins involved in the ubiquitin cascade) were expressed in *Escherichia coli* BL21 (DE3) cells. Cells were grown at 37°C in Luria-Bertani (LB) medium to an OD_600_ of approximately 0.6, then induced with 0.2 mM isopropyl-β-D-1-thiogalactopyranoside (IPTG) and grown at 18°C overnight. Cell pellets were resuspended in purification buffer [50 mM Tris (pH 8.0), 300 mM NaCl, 10% (v/v) glycerol, and 10% (w/v) sucrose] supplemented with lysozyme (0.2 mg/ml). Cells were lysed by sonication and then purified by affinity purification, as outlined below. Ubiquitin was expressed as an untagged protein and then purified and labelled with Cy3 as per ref^53^.TRIB1 C-terminus (345-372) was first purified using Ni^2+^ affinity resin (His-Select resin (Sigma-Aldrich) before being dialysed into PBS pH 8.0. The Alexa Fluor™ 647 Protein Labelling Kit (A20173 (ThermoFisher)) was used to label 2.7 mg TRIB1. Labelled protein was then snap frozen in liquid nitrogen and stored at -80°C.

The DDD complex was produced in either *Trichoplusia ni* (TNi) (Full-length DDB1, DET1, DDA1) or *Spodoptera frugiperda* (SF9) at 27°C (Full-length DDB1, DET1, DDA1 and Ube2e2) or 22°C (DDD(ΔBPB), Full-length DET1 and DDA1 (DDD(ΔBPB))). Full-length COP1 was produced in TNi cells. Cells were incubated with baculovirus at 28°C for 72 hours (Sf9) or 48 hours (TNi) or at 22°C for 96 hours (Sf9). Cells were harvested by centrifugation at 1000 xg for 5 min. Cell pellets were resuspended in 50 mM Tris-HCl (pH 8.0), 10% (v/v) glycerol, 150 mM NaCl, and 10 mM imidazole. Cells were lysed via sonication, and the soluble fraction was purified by affinity chromatography (either Ni^2+^ or Strep) (His-Select resin (Sigma-Aldrich); Strep-Tactin®XT 4Flow resin (IBA Lifesciences)). Affinity purification was performed by incubating clarified lysates with resin, washing with an appropriate buffer and then eluting the protein from the resin with either a buffer containing biotin (Buffer BXT) for proteins bound to Strep-Tactin®XT 4Flow resin, or with a buffer containing 300 mM imidazole for those bound to His-Select resin. COP1 and the DDD complexes were purified using StrepTactin®XT 4Flow resin. All other recombinant proteins were purified using Ni^2+^ affinity purification, unless otherwise indicated. If further purification was required, this was done by size exclusion chromatography using either a Superdex200 Increase column (GE Life Sciences) [150-300 mM NaCl and 50 mM Tris-HCl buffer (pH 8), 2 mM DTT or 50 mM Tricine pH 8, 80 mM NaCl, 0.5 mM TCEP], or a Superose 6 Increase Column (GE Life Sciences). Purified protein was snap frozen and stored at –80°C until required. The full-length DDD required for multi-turnover assays was purified in PBS.

If required for discharge assays, following size exclusion fractions containing the DDD(ΔBPB) complex were pooled and dialysed into 50 mM Tris pH 8, 150 mM NaCl If required for cryoEM the DDD complex was purified with a buffer containing more NaCl and in the absence of a reducing agent. The initial full-length DDD complex for EM was purified in the aforementioned Tricine Buffer but all further purifications used a Tris Buffer.

### Electron Microscopy and Structure determination

For full-length DDB1-DET1-DDA1-Ube2e2, glow-discharged 300-mesh C-Flat 1.2 / 1.3 grids (Protochips) were placed in Vitrobot MarkIV (ThermoFisher Scientific) at 4°C and 100% relative humidity. 3 µL of protein sample at 0.6 mg/ml concentration in 50 mM Tricine pH 8, 80 mM NaCl, 0.5 mM TCEP was applied to the grids and blotted for 8 seconds with blot force 4 before plunge-freezing in liquid ethane. 4957 micrographs were collected using a Titan Krios microscope (Thermo Fisher Scientific) operated at 300 kV. Micrographs were recorded using EPU with a K2 summit (Gatan) camera at calibrated pixel size 0.66 Å /px and 0.8-2 μm underfocus. The total dose of 66 e / Å ^2^ was fractionated over 60 frames. Processing was carried out in Cryosparc v 3.0.1, unless specified otherwise^50^. Movies were aligned patchwise and binned 2x2. CTFs of the aligned micrographs were determined patchwise.

Blob-picking and 2D classification yielded templates for template-based picking, where further 2D classification and selection gave a dataset of 550k particles. Rounds of 2D and 3D classification yielded a set of 221k particles which were refined in non-uniform refinement to 3 Å resolution^51^. The micrographs were motion-corrected in a full-frame manner and full-resolution particles with per-particle, local motion correction were then reconstructed in a non-uniform refinement to a resolution of 2.89 Å —but which suffered from severe map anisotropy. Processing in parallel using 3D classifications, multi-body refinements, or focused classifications and refinements in RELION 3.1.2 ^52,53^ did not improve map clarity. The final map from cryosparc was post-processed using deepEMhancer^54^, and its sphericity, global and directional resolution was estimated using 3DFSC^55^.

The DDD(ΔBPB) complex was concentrated, mixed with an excess of Ube2e2 and then re-purified using an S200 Increase 5/150 in 50 mM Tris pH 8, 300 mM NaCl. Fractions containing the whole complex were pooled and then concentrated to ∼0.85 mg/mL before cryoEM. Samples for collection were placed on Quantifoil 1.2/1.3 grids in a Vitrobot MarkIV (ThermoFisher Scientific) at 10 degrees C and 100% relative humidity. 3 µL of protein sample was applied to the grids and blotted for 5 seconds before plunge-freezing in liquid ethane. 14499 micrographs were collected using a Titan Krios microscope (Thermo Fisher Scientific) operated at 300 kV. Micrographs were recorded using EPU equipped with a BioQuantum energy filter (Gatan) (15 eV) and a K3 direct detection camera (Gatan) at calibrated pixel size 0.65 Å /px and -1.0 μm underfocus. The total dose of 80 e / Å ^2^ was fractionated over 100 frames. Processing was carried out mainly in Cryosparc v4.3.1, with some later processing using versions up to v4.4.1+. Movies and CTFs were aligned patchwise. Blob picking and 2D classification yielded templates for template-based picking, where further 2D classification and selection yielded 1.9 million particles. Rounds of 3D classification yielded a set of 375k particles, which were then further divided through 3D variability analysis using a mask of the whole complex. This yielded a set of 63k particles, which were refined in non-uniform refinement to 3 Å resolution. The micrographs of those particles were re-extracted at a larger box size before local and global CTF refinement was performed. Reference based motion correction was performed on the exposures that yielded those particles before a final non-uniform refinement was performed, resulting in a set of 61,234 particles yielding a resolution of 2.83 Å. Structural determination was performed iteratively using Phenix and Coot to fit the structural model to the map refined in Cryosparc.

### AlphaFold Modelling

Alphafold2 modelling was performed on the high-performance computing cluster at the University of Otago. Models of DET1-COP1 binding were generated using AlphaFold2-Multimer^54,55^ based on full HHblits alignment protocols implemented in https://github.com/google-deepmind/alphafold, with max_template_date=2020-05-14, model_preset=multimer and 5 predictions per model. The N-terminal 125 amino acids of COP1, which are suggested to be completely disordered in full-length AlphaFold models were omitted from predictions. Models of E2 enzymes in complex with DET1 were generated using full-length DET1, the full-length or full-length human sequences of Ube2e2, or core catalytic domains of other E2 enzymes as displayed in the alignment in Supplementary Figure 4. Top models of COP1-DET1, or DET1-E2s were ranked based on iPTM score, and highest scoring models, or summary data corresponding to iPTM scores, are represented in the manuscript.

### Immunoprecipitation

A series of Ubiquitin-conjugating enzymes (E2) were expressed in *Escherichia coli* BL21 (DE3) cells. All constructs contained a His_6_ affinity tag, either N-terminal (Ubc13, Ube2W, Ube2e) or C-terminal (UbeE2K, Ube2T (1-153) Ube2T_FL, UbcH5b). Cells were grown at 37°C in Luria-Bertani (LB) medium to an OD_600_ of approximately 0.6, then induced with 0.2 mM isopropyl-β-D-1-thiogalactopyranoside (IPTG) and grown at 18°C overnight. Cells were lysed by sonication and the proteins purified by Ni^2+^ affinity chromatography as described above. Meanwhile the complex containing full-length DDB1, DET1 and DDA1 was produced in TNi cells. The complex was immobilised on Strep-Tactin®XT 4Flow resin (IBA Lifesciences) and analysed for purity by SDS-PAGE. The His_6_-E2 proteins were incubated with either the DDD resin or Strep-Tactin®XT 4Flow resin which was unbound at 4°C for 20 min with Buffer W with 0.02% (v/v) Tween 20 and 2 mM DTT. After incubation, the resins were washed three times in this buffer. Samples were then resuspended in SDS sample buffer and visualized on 10–20% gradient SDS-PAGE stained with Coomassie R250.

### Ubiquitination Assays

Multi-turnover activity assays were carried out at 37°C for the indicated times to determine E3 activity of COP1. Reactions were carried out in a buffer containing 50 mM Tris pH 7.5, 50 mM NaCl, 2 mM MgCl_2_, 2 mM TCEP pH 7.0, 5 mM ATP pH 7.0. Assays were carried out containing E1 (0.1 μM), E2 (10 μM), and Cy3-labelled ubiquitin (50–100 μM) (final concentration). Either UbcH5b or Ube2e2 were used as E2. When these were combined, each was used at 10 μM. Samples were incubated at 37°C for the indicated time points and the reaction stopped by the addition of 4 × SDS–PAGE sample buffer. Cy3-labelled ubiquitin was detected using the Odyssey Fc imaging system (600 nM) prior to staining the gel with Coomassie R250.

Ube2e2 Charging: a 100 µL reaction mix containing E1 (1 µM), dN-Ube2e2 (60 µM), and Ubiquitin (150 µM) was incubated at 37°C for 5 minutes to charge Ube2e2. Reactions were carried out in a buffer containing 50 mM Tris pH 7.5, 50 mM NaCl, 2 mM MgCl_2_, 2 mM TCEP pH 7.0, 5 mM ATP pH 7 .0. Following charging, the reaction mix was loaded onto an Superdex 75 Increase column (GE Life Sciences) and separated using a buffer containing 50 mM Tris pH 8, 150 mM NaCl. The resulting fractions which contained charged Ube2e2 were collected and used in discharge assays, as outlined below.

Discharge assays were performed at 37°C for the indicated times to determine whether the DDD complex possessed any ability to cause ubiquitin discharge from charged E2 (Ube2e2). Reactions were performed in a buffer containing 50 mM Tris pH 8, 150 mM NaCl. Assays contained Ube2e2-Ub conjugate and purified DDD(ΔBPB) complex (1 µM). The positive and negative controls for these assays either replaced the DDD(ΔBPB) complex with RNF12 or omitted the DDD(ΔBPB) complex, respectively. Reactions were quenched with non-reducing SDS sample buffer and analysed via SDS-PAGE stained with Coomassie R250.

### Cell lines and cell culture

Expi293F were grown in Expi293 Expression Medium (Gibco, A1435102) supplemented with Penicillin-Streptomycin (100 U/ml; Life Technologies, 15140122). Cells were grown at 37°C in a humidified atmosphere with 8% CO_2_, shaking at 120 rpm. Expi293F cells were transiently co-transfected with Polyethylenimine (PEI 25K, Australia Bioscientific. PEI MAX-Transfection Grade Linear Polyethylenimine Hydrochloride, 24765-1) to express either COP1(wild-type/mutants) or DET1(full-length/Δcup). Cells were grown for 72 hours before being harvested by centrifugation at 800 xg, 10 mins, 4°C. The cell pellet was washed with cold dPBS and then the cells incubated with RIPA buffer for 30 mins on ice to lyse. The lysate was clarified by centrifugation at 10,000 xg, 4°C 10 min and the supernatant snap frozen in liquid nitrogen, to be used for subsequent analysis. All protein lysates were quantified using a BCA assay (Pierce™ BCA Protein Assay) and analysed by western blotting.

### Co-Purification of DET1 by COP1

COP1 (both wild-type and loop mutants) and DET1 (full-length/cup deletion mutant, Δ) were expressed separately in Expi293F. Cells were grown for 72 hours before being harvested by centrifugation at 800 xg, 10 mins, 4°C. The pellets of these were mixed before lysis by sonication and affinity purification of the StrepII-COP1. Clarified lysates were incubated with StrepTactin ®XT 4Flow resin for 20 mins at 4°C in rotation. The resin was washed four times with Buffer W supplemented with 0.02 % (v/v) Tween 20 then resuspended in 4 × SDS–PAGE sample buffer and then western blotted to determine if DET1 could be retained by COP1. Lysates containing only DET1 were bound to StrepTactin®XT 4Flow resin to access non-specific binding of DET1 to the resin itself.

### Western blotting

For analysis by western blot, samples were separated by SDS/PAGE and transferred to 0.2 μm nitrocellulose (Life Technologies, IB23002). Membranes were blocked in 5% BSA (w/v) in TBS-T. Membranes were incubated with primary antibodies overnight at 4°C in 1% BSA (w/v) in TBS-T. Antibodies used were: Rabbit α-HA (1:5000, CST 3724S), mouse α-COP1 (1:1000, Santa Cruz, C-terminal epitope, sc166799) and/or mouse Anti-α -tubulin (clone DM1A) (1:10,000, Millipore).

Following three washes with TBS-T, membranes were incubated with secondary antibodies diluted in TBS-T with 1% (w/v) BSA for 1 hour at room temperature. Secondary antibodies used were goat anti-rabbit IRdye 680LT (LI-COR) or goat anti-mouse IRdye 800LT (LI-COR). Membranes were washed a further three times with TBS-T. Membranes were then developed with the Odyssey Fc imaging system.

## Reference

1. Padovani, C., Jevtić, P., and Rapé, M. (2022). Quality control of protein complex composition. Mol. Cell 0. 10.1016/j.molcel.2022.02.029.

2. Swatek, K.N., and Komander, D. (2016). Ubiquitin modifications. Cell Res. 26, 399–422.

3. Lau, O.S., and Deng, X.-W. (2012). The photomorphogenic repressors COP1 and DET1: 20 years later. Trends Plant Sci. 17, 584–593.

4. Hoecker, U. (2017). The activities of the E3 ubiquitin ligase COP1/SPA, a key repressor in light signaling. Curr. Opin. Plant Biol. 37, 63–69.

5. Ponnu, J., and Hoecker, U. (2021). Illuminating the COP1/SPA Ubiquitin Ligase: Fresh Insights Into Its Structure and Functions During Plant Photomorphogenesis. Front. Plant Sci. 12, 662793.

6. Dedhia, P.H., Keeshan, K., Uljon, S., Xu, L., Vega, M.E., Shestova, O., Zaks-Zilberman, M., Romany, C., Blacklow, S.C., and Pear, W.S. (2010). Differential ability of Tribbles family members to promote degradation of C/EBPalpha and induce acute myelogenous leukemia. Blood 116, 1321–1328.

7. Jamieson, S.A., Ruan, Z., Burgess, A.E., Curry, J.R., McMillan, H.D., Brewster, J.L., Dunbier, A.K., Axtman, A.D., Kannan, N., and Mace, P.D. (2018). Substrate binding allosterically relieves autoinhibition of the pseudokinase TRIB1. Sci. Signal. 11. 10.1126/scisignal.aau0597.

8. Vitari, A.C., Leong, K.G., Newton, K., Yee, C., O’Rourke, K., Liu, J., Phu, L., Vij, R., Ferrando, R., Couto, S.S., et al. (2011). COP1 is a tumour suppressor that causes degradation of ETS transcription factors. Nature 474, 403–406.

9. Wertz, I.E., O’Rourke, K.M., Zhang, Z., Dornan, D., Arnott, D., Deshaies, R.J., and Dixit, V.M. (2004). Human De-etiolated-1 regulates c-Jun by assembling a CUL4A ubiquitin ligase. Science 303, 1371–1374.

10. Zhang, Z., Newton, K., Kummerfeld, S.K., Webster, J., Kirkpatrick, D.S., Phu, L., Eastham-Anderson, J., Liu, J., Lee, W.P., Wu, J., et al. (2017). Transcription factor Etv5 is essential for the maintenance of alveolar type II cells. Proc. Natl. Acad. Sci. U. S. A. 114, 3903–3908.

11. Bauer, R.C., Sasaki, M., Cohen, D.M., Cui, J., Smith, M.A., Yenilmez, B.O., Steger, D.J., and Rader, D.J. (2015). Tribbles-1 regulates hepatic lipogenesis through posttranscriptional regulation of C/EBPα. J. Clin. Invest. 125, 3809–3818.

12. Suriben, R., Kaihara, K.A., Paolino, M., Reichelt, M., Kummerfeld, S.K., Modrusan, Z., Dugger, D.L., Newton, K., Sagolla, M., Webster, J.D., et al. (2015). β-Cell Insulin Secretion Requires the Ubiquitin Ligase COP1. Cell 163, 1457–1467.

13. Wang, X., Tokheim, C., Gu, S.S., Wang, B., Tang, Q., Li, Y., Traugh, N., Zeng, Z., Zhang, Y., Li, Z., et al. (2021). In vivo CRISPR screens identify the E3 ligase Cop1 as a modulator of macrophage infiltration and cancer immunotherapy target. Cell 184, 5357–5374.e22.

14. Marine, J.-C. (2012). Spotlight on the role of COP1 in tumorigenesis. Nat. Rev. Cancer 12, 455–464.

15. Ndoja, A., Reja, R., Lee, S.-H., Webster, J.D., Ngu, H., Rose, C.M., Kirkpatrick, D.S., Modrusan, Z., Chen, Y.-J.J., Dugger, D.L., et al. (2020). Ubiquitin Ligase COP1 Suppresses Neuroinflammation by Degrading c/EBPβ in Microglia. Cell 182, 1156–1169.e12.

16. Xie, Y., Cao, Z., Wong, E.W., Guan, Y., Ma, W., Zhang, J.Q., Walczak, E.G., Murphy, D., Ran, L., Sirota, I., et al. (2018). COP1/DET1/ETS axis regulates ERK transcriptome and sensitivity to MAPK inhibitors. J. Clin. Invest. 128, 1442–1457.

17. Nixdorf, M., and Hoecker, U. (2010). SPA1 and DET1 act together to control photomorphogenesis throughout plant development. Planta 231, 825–833.

18. Scrima, A., Koníčková, R., Czyzewski, B.K., Kawasaki, Y., Jeffrey, P.D., Groisman, R., Nakatani, Y., Iwai, S., Pavletich, N.P., and Thomä, N.H. (2008). Structural basis of UV DNA-damage recognition by the DDB1-DDB2 complex. Cell 135, 1213–1223.

19. Mohamed, W.I., Schenk, A.D., Kempf, G., Cavadini, S., Basters, A., Potenza, A., Abdul Rahman, W., Rabl, J., Reichermeier, K., and Thomä, N.H. (2021). The CRL4DCAF1 cullin-RING ubiquitin ligase is activated following a switch in oligomerization state. EMBO J. 40, e108008.

20. Fonseca, S., and Rubio, V. (2019). Arabidopsis CRL4 Complexes: Surveying Chromatin States and Gene Expression. Front. Plant Sci. 10, 1095.

21. Petroski, M.D., and Deshaies, R.J. (2004). Function and regulation of cullin-RING ubiquitin ligases. Nat. Rev. Mol. Cell Biol. 6, 9–20.

22. Li, T., Robert, E.I., van Breugel, P.C., Strubin, M., and Zheng, N. (2010). A promiscuous alpha-helical motif anchors viral hijackers and substrate receptors to the CUL4-DDB1 ubiquitin ligase machinery. Nat. Struct. Mol. Biol. 17, 105–111.

23. Baek, K., Scott, D.C., and Schulman, B.A. (2021). NEDD8 and ubiquitin ligation by cullin-RING E3 ligases. Curr. Opin. Struct. Biol. 67, 101–109.

24. Wei, N., and Deng, X.-W. (2003). The COP9 signalosome. Annu. Rev. Cell Dev. Biol. 19, 261–286.

25. Cavadini, S., Fischer, E.S., Bunker, R.D., Potenza, A., Lingaraju, G.M., Goldie, K.N., Mohamed, W.I., Faty, M., Petzold, G., Beckwith, R.E.J., et al. (2016). Cullin-RING ubiquitin E3 ligase regulation by the COP9 signalosome. Nature 531, 598–603.

26. Békés, M., Langley, D.R., and Crews, C.M. (2022). PROTAC targeted protein degraders: the past is prologue. Nat. Rev. Drug Discov., 1–20.

27. Shabek, N., Ruble, J., Waston, C.J., Garbutt, K.C., Hinds, T.R., Li, T., and Zheng, N. (2018). Structural insights into DDA1 function as a core component of the CRL4-DDB1 ubiquitin ligase. Cell Discov 4, 67.

28. Pick, E., Lau, O.S., Tsuge, T., Menon, S., Tong, Y., Dohmae, N., Plafker, S.M., Deng, X.-W., and Wei, N. (2007). Mammalian DET1 regulates Cul4A activity and forms stable complexes with E2 ubiquitin-conjugating enzymes. Mol. Cell. Biol. 27, 4708– 4719.

29. Suzuki, G., Yanagawa, Y., Kwok, S.F., Matsui, M., and Deng, X.-W. (2002). Arabidopsis COP10 is a ubiquitin-conjugating enzyme variant that acts together with COP1 and the COP9 signalosome in repressing photomorphogenesis. Genes Dev. 16, 554–559.

30. Yanagawa, Y., Sullivan, J.A., Komatsu, S., Gusmaroli, G., Suzuki, G., Yin, J., Ishibashi, T., Saijo, Y., Rubio, V., Kimura, S., et al. (2004). Arabidopsis COP10 forms a complex with DDB1 and DET1 in vivo and enhances the activity of ubiquitin conjugating enzymes. Genes Dev. 18, 2172–2181.

31. Uljon, S., Xu, X., Durzynska, I., Stein, S., Adelmant, G., Marto, J.A., Pear, W.S., and Blacklow, S.C. (2016). Structural Basis for Substrate Selectivity of the E3 Ligase COP1. Structure 24, 687–696.

32. Angers, S., Li, T., Yi, X., MacCoss, M.J., Moon, R.T., and Zheng, N. (2006). Molecular architecture and assembly of the DDB1-CUL4A ubiquitin ligase machinery. Nature 443, 590–593.

33. Watson, E.R., Novick, S., Matyskiela, M.E., Chamberlain, P.P., H de la Peña, A., Zhu, J., Tran, E., Griffin, P.R., Wertz, I.E., and Lander, G.C. (2022). Molecular glue CELMoD compounds are regulators of cereblon conformation. Science 378, 549–553.

34. Bussiere, D.E., Xie, L., Srinivas, H., Shu, W., Burke, A., Be, C., Zhao, J., Godbole, A., King, D., Karki, R.G., et al. (2020). Structural basis of indisulam-mediated RBM39 recruitment to DCAF15 E3 ligase complex. Nat. Chem. Biol. 16, 15–23.

35. Du, X., Volkov, O.A., Czerwinski, R.M., Tan, H., Huerta, C., Morton, E.R., Rizzi, J.P., Wehn, P.M., Xu, R., Nijhawan, D., et al. (2019). Structural Basis and Kinetic Pathway of RBM39 Recruitment to DCAF15 by a Sulfonamide Molecular Glue E7820. Structure 27, 1625–1633.e3.

36. Brzovic, P.S., Lissounov, A., Christensen, D.E., Hoyt, D.W., and Klevit, R.E. (2006). A UbcH5/ubiquitin noncovalent complex is required for processive BRCA1-directed ubiquitination. Mol. Cell 21, 873–880.

37. Stewart, M.D., Ritterhoff, T., Klevit, R.E., and Brzovic, P.S. (2016). E2 enzymes: more than just middle men. Cell Res. 26, 423–440.

38. Plechanovova, A., Jaffray, E.G., Tatham, M.H., Naismith, J.H., and Hay, R.T. (2012). Structure of a RING E3 ligase and ubiquitin-loaded E2 primed for catalysis. Nature 489, 115–120.

39. Buetow, L., Gabrielsen, M., Anthony, N.G., Dou, H., Patel, A., Aitkenhead, H., Sibbet, G.J., Smith, B.O., and Huang, D.T. (2015). Activation of a primed RING E3-E2-ubiquitin complex by non-covalent ubiquitin. Mol. Cell 58, 297–310.

40. Middleton, A.J., Zhu, J., and Day, C.L. (2020). The RING domain of RING finger 12 efficiently builds degradative ubiquitin chains. J. Mol. Biol. 432, 3790–3801.

41. Abramson, J., Adler, J., Dunger, J., Evans, R., Green, T., Pritzel, A., Ronneberger, O., Willmore, L., Ballard, A.J., Bambrick, J., et al. (2024). Accurate structure prediction of biomolecular interactions with AlphaFold 3. Nature. 10.1038/s41586-024-07487-w.

42. Baek, K., Krist, D.T., Prabu, J.R., Hill, S., Klügel, M., Neumaier, L.-M., von Gronau, S., Kleiger, G., and Schulman, B.A. (2020). NEDD8 nucleates a multivalent cullin-RING-UBE2D ubiquitin ligation assembly. Nature 578, 461–466.

43. Christensen, D.E., Brzovic, P.S., and Klevit, R.E. (2007). E2-BRCA1 RING interactions dictate synthesis of mono- or specific polyubiquitin chain linkages. Nat. Struct. Mol. Biol. 14, 941–948.

44. Banka, P.A., Behera, A.P., Sarkar, S., and Datta, A.B. (2015). RING E3-Catalyzed E2 Self-Ubiquitination Attenuates the Activity of Ube2E Ubiquitin-Conjugating Enzymes. J. Mol. Biol. 427, 2290–2304.

45. Nguyen, L., Plafker, K.S., Starnes, A., Cook, M., Klevit, R.E., and Plafker, S.M. (2014). The ubiquitin-conjugating enzyme, UbcM2, is restricted to monoubiquitylation by a two-fold mechanism that involves backside residues of E2 and Lys48 of ubiquitin. Biochemistry 53, 4004–4014.

46. Schumacher, F.-R., Wilson, G., and Day, C.L. (2013). The N-terminal extension of UBE2E ubiquitin-conjugating enzymes limits chain assembly. J. Mol. Biol. 425, 4099– 4111.

47. Li, Y.-D., Ma, M.W., Hassan, M.M., Hunkeler, M., Teng, M., Puvar, K., Lumpkin, R., Sandoval, B., Jin, C.Y., Ficarro, S.B., et al. (2023). Template-assisted covalent modification of DCAF16 underlies activity of BRD4 molecular glue degraders. bioRxiv, 2023.02.14.528208. 10.1101/2023.02.14.528208.

48. Hsia, O., Hinterndorfer, M., Cowan, A.D., Iso, K., Ishida, T., Sundaramoorthy, R., Nakasone, M.A., Imrichova, H., Schätz, C., Rukavina, A., et al. (2024). Targeted protein degradation via intramolecular bivalent glues. Nature 627, 204–211.

49. Eyers, P.A., Keeshan, K., and Kannan, N. (2017). Tribbles in the 21st Century: The Evolving Roles of Tribbles Pseudokinases in Biology and Disease. Trends Cell Biol. 27, 284–298.

50. Jamieson, S.A., Pudjihartono, M., Horne, C.R., Viloria, J.S., Dunlop, J.L., McMillan, H.D., Day, R.C., Keeshan, K., Murphy, J.M., and Mace, P.D. (2022). Nanobodies identify an activated state of the TRIB2 pseudokinase. Structure 30, 1518–1529.e5.

51. Reichermeier, K.M., Straube, R., Reitsma, J.M., Sweredoski, M.J., Rose, C.M., Moradian, A., den Besten, W., Hinkle, T., Verschueren, E., Petzold, G., et al. (2020). PIKES Analysis Reveals Response to Degraders and Key Regulatory Mechanisms of the CRL4 Network. Mol. Cell 77, 1092–1106.e9.

52. Selvaraj, N., Kedage, V., and Hollenhorst, P.C. (2015). Comparison of MAPK specificity across the ETS transcription factor family identifies a high-affinity ERK interaction required for ERG function in prostate cells. Cell Commun. Signal. 13, 12.

53. Paluda, A., Middleton, A.J., Rossig, C., Mace, P.D., and Day, C.L. (2022). Ubiquitin and a charged loop regulate the ubiquitin E3 ligase activity of Ark2C. Nat. Commun. 13, 1181.

54. Evans, R., O’Neill, M., Pritzel, A., Antropova, N., Senior, A., Green, T., Žídek, A., Bates, R., Blackwell, S., Yim, J., et al. (2022). Protein complex prediction with AlphaFold-Multimer. bioRxiv, 2021.10.04.463034. 10.1101/2021.10.04.463034.

55. Jumper, J., Evans, R., Pritzel, A., Green, T., Figurnov, M., Ronneberger, O., Tunyasuvunakool, K., Bates, R., Žídek, A., Potapenko, A., et al. (2021). Highly accurate protein structure prediction with AlphaFold. Nature 596, 583–589.

